# Tumor-educated Gr1^+^CD11b^+^ cells instigate breast cancer metastasis by twisting cancer cells plasticity via OSM/IL6–JAK signaling

**DOI:** 10.1101/2022.10.26.513889

**Authors:** Sanam Peyvandi, Manon Bulliard, Annamaria Kauzlaric, Oriana Coquoz, Yu-Ting Huang, Nathalie Duffey, Laetitia Gafner, Girieca Lorusso, Nadine Fournier, Qiang Lan, Curzio Rüegg

## Abstract

Cancer cell plasticity contributes to tumor therapy resistance and metastasis formation, which represent the main causes of cancer-related death for most cancers, including breast cancer. The tumor microenvironment drives cancer cell plasticity and metastasis and, thus, unravelling the underlying cues may provide novel effective strategies to manage metastatic disease. Here, we show that stem cell antigen-1 positive (Sca-1^+^) murine breast cancer cells enriched during tumor progression and metastasis have higher *in vitro* cancer stem cell-like properties, enhanced *in vivo* metastatic ability, and initiate primary tumors rich in Gr1^high^CD11b^+^Ly6C^low^ cells. In turn, tumor-educated Gr1^+^CD11b^+^ (Tu-Gr1^+^CD11b^+^) cells rapidly and transiently convert low metastatic 4T1-Sca-1^-^ cells into highly metastatic 4T1-Sca-1^+^ cells via secreted OSM and IL6. Moreover, chemotherapy-resistant and highly metastatic 4T1-derived cells maintain high Sca-1^+^ frequency through cell autonomous IL6 production. Inhibition of OSM, IL6 or JAK suppressed Tu-Gr1^+^CD11b^+^-induced Sca-1^+^ population enrichment *in vitro*, while JAK inhibition abrogated metastasis of chemotherapy-enriched Sca-1^+^ cells *in vivo*. Importantly, Tu-Gr1^+^CD11b^+^ cells invoked a gene signature in tumor cells predicting shorter OS and RFS in breast cancer patients. Collectively, our data identified OSM/IL6-JAK as a clinically relevant paracrine/autocrine axis instigating breast cancer cell plasticity triggering metastasis.

## Introduction

Metastasis accounts for over 90% of cancer-related death, calling for new strategies to prevent cancer cell dissemination and metastasis formation (1). Recent studies using single-cell lineage tracing and single-cell RNA sequencing (scRNA-seq) technologies have provided detailed information about intratumor heterogeneity (2, 3), whereby genetically, epigenetically and functionally diverse subpopulations of cancer cells exist within the tumor mass, spatially and temporally (4). Intratumor heterogeneity may arise by modulating cancer cell plasticity, especially of cancer stem cells (CSCs), through cell-intrinsic and - extrinsic mechanisms (5, 6).

Multiple CSC subpopulations appear to co-exist within the primary tumor mass resulting in high degree of tumor cell heterogeneity and increased aggressiveness (7) including in breast cancer (4, 8–11). In particular, CSCs can acquire metastasis initiating capacities (12) and resistance to therapy, resulting in cancer relapse (4). Moreover, non-CSCs within the tumor bulk may acquire CSC properties to repopulate the tumor(4). While CSC are defined rather functionally by their ability to initiate tumors and metastasis in low number *in vivo*, several cell surface markers associated with CSC features have been reported, including CD44, CD24, Sca-1, CD61, CD49f were used to identify breast CSCs (13, 14).

The interaction of tumor cells with the tumor microenvironment (TME) contributes to tumor cell plasticity and tumor heterogeneity (15, 16). Cells of the TME also promote tumor escape and metastasis through multiple mechanisms, including promotion of angiogenesis, cell survival, invasion, epithelial-mesenchymal transition (EMT), and immunosuppression (17–22). Recently, they have been reported to instigate expansion of CSC with metastatic ability (also known as metastasis-initiating cells) in different cancers (17, 23, 24). Thus, the TME dynamics is a key driver of cancer cell plasticity and heterogeneity promoting tumor growth, progression and metastasis (4). Accurate characterization of the regulation of tumor plasticity and heterogeneity by the TME may reveal novel opportunities for developing effective anti-metastatic therapies (8, 14).

TME-derived Oncostatin M (OSM) has been shown to mediate tumor progression and CSC stemness by activating its receptor OSMR (25). OSM belongs to the IL6 family of cytokines (including IL6 itself, IL11 and LIF) (26, 27), whose members bind to dimeric receptors sharing a common subunit (gp130 or IL6ST) and activate JAK-STAT, RAS-MAPK and PI3K-AKT pathways (28, 29). Increased OSM or IL6 expression correlates with reduced survival in breast cancer patients (30, 31). OSM was shown to drive breast cancer progression and metastasis through direct effects on cancer cells, such as suppression of estrogen receptor (ER) expression (31) and promotion of EMT (25, 32), and indirect effects via TME cells, in particular the reprogramming of tumor associated macrophages and fibroblasts (33–36).

Here, by assessing the metastatic evolution of murine triple-negative breast cancer (TNBC) models *in silico* and *in vivo*, we observed that the Sca-1^+^ tumor cell subpopulation is enriched during tumor progression. We show that tumor-educated Gr1^+^CD11b^+^ cells (Tu-Gr1^+^CD11b^+^), but not naïve Gr1^+^CD11b^+^ cells from spleen (Spl-Gr1^+^CD11b^+^) or bone marrow (BM-Gr1^+^CD11b^+^) in tumor-bearing mice, modulate tumor plasticity via OSM/IL6-JAK signaling by rapidly and transiently converting 4T1-Sca-1^-^ cells into 4T1-Sca-1^+^ cells with high metastatic capacity. Prolonged exposure of 4T1 cells to chemotherapy stably enriched for metastatic Sca-1^+^ cells via an autocrine IL6-JAK signaling loop. A short *in vitr*o treatment of these chemo-resistant cells with the JAK inhibitor Ruxolitinib, suppressed their metastatic capacity. Importantly, Tu-Gr1^+^CD11b^+^ invoked a gene expression signature in 4T1 cells that predicted shorter overall survival (OS) and relapse-free survival (RFS) in breast cancer patients, reinforcing the clinical relevance of these findings.

Our results reveal a novel mechanism modulating tumor plasticity and triggering the emergence of cancer cells with enhanced metastatic capacity, through paracrine (Tu-Gr1^+^CD11b^+^-mediated) and cell autonomous (chemotherapy-induced) OSM/IL6-JAK dependent signaling. The OSM/IL6-JAK axis may be considered as a candidate of actionable clinical targets to impinge on metastatic progression and therapy resistance.

## Results

### A Sca-1^+^ tumor cell population is enriched during tumor progression and has higher *in vivo* metastatic capacity

To investigate tumor cell heterogeneity during tumor progression, we first examined the expression of the previously reported breast CSC markers CD24, CD44, CD61, Sca-1 and CD49f (13) in the publicly available RNA sequencing (RNAseq) dataset from Ross *et al.* encompassing several murine breast cancer models (37). This dataset includes data from cultured cancer cells (In_Culture), orthotopic primary tumors (OT_PT), spontaneous lung metastases (OT_LuM), and experimental lung metastases after tail vein injection (TV_LuM) (Figure 1A and Supplemental Figure 1A). *Sca-1* expression was elevated in lung metastasis in 4T1, 6DT1, Mvt1 and Met1 models compared with the respective primary tumors. Interestingly, in 4T1, 6DT1 and Mvt1 models, *Sca-1* expression was also elevated in experimental lung metastases compared with cultured cells. However, the expression of *Cd24*, *Cd44*, *Cd61* and *Cd49f* were not altered, or their expression pattern was inconsistent during progression (Supplemental Figure 1A). Thus, increased *Sca-1* expression during metastasis is consistently observed in different preclinical breast cancer models.

**Figure 1.**
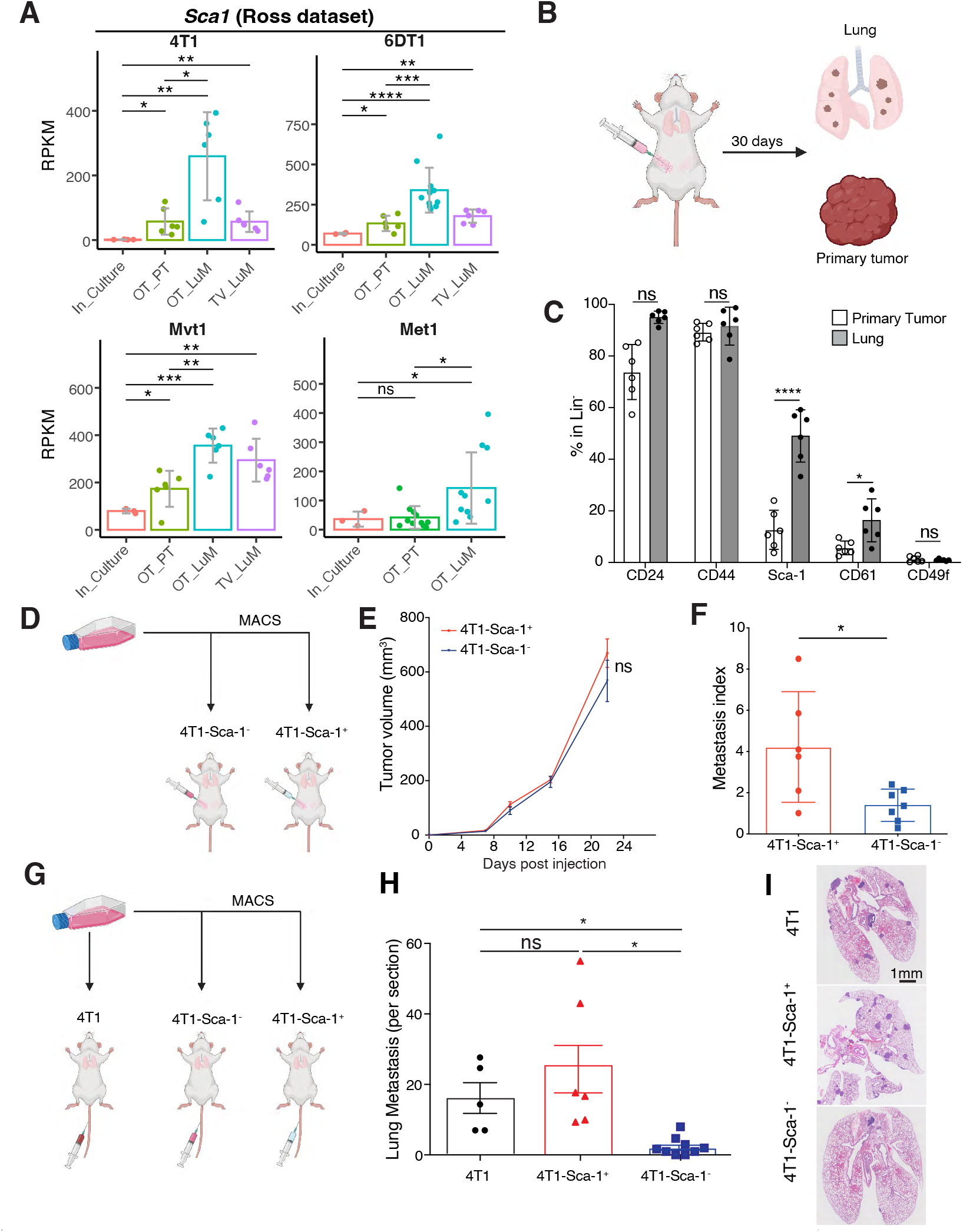
Sca-1+ population is enriched during in vivo metastasis across multiple breast cancer models. (**A**) *Sca-1* mRNA expression in the metastatic murine breast cancer models 4T1, 6DT1, Mvt1 and Met1, extracted from the Ross dataset. Analyzed samples consist of cultured cells (In_Culture), orthotopic injected primary tumors (OP_PT), spontaneous lung metastases (OP_LuM) and lung metastases induced by tail-vain injection (TV_LuM). Data are presented as mean of Reads Per Kilobase of transcript per Million mapped reads (RPKM) ± SD (unpaired two-tailed student’s t test with Holm correction). (**B**) Experimental set up for *in vivo* experimental validation. The 4T1 tumor cells were orthotopically injected into the 4th mammary fat pad. Thirty days later, cells from primary tumors and lungs were isolated to examine CSC marker expression by flow cytometry. (**C**) Frequency of CSC marker expression in primary tumors and lung metastases. Results give the percentage of CD24, CD44, Sca-1, CD61 and CD49f positive cells gated in lineage negative cells (CD45-CD31-TER119-). (**D-F**) Experimental set up (**D**) of the in vivo experiment to assess tumor growth (e) and lung metastatic ability (metastatic index) (**F**) of 4T1-Sca-1+ and 4T1-Sca-1- populations isolated from parental 4T1 cells orthotopically injected into the 4th mammary fat pad. Metastases are assessed 21 days after tumor cell injection. n=8/group, 3 independent experiments. (**G-I**) Experimental set up (**G**) of the in vivo experiment to assess lung colonization capacity of sorted parental 4T1, 4T1-Sca-1+ and 4T1-Sca-1- cells by tail-vain injection. Lung metastatic nodule numbers (**H**) and representative images (**I**) of lungs from mice 10 days post injection (n=5-6, 2 independent experiments). Scale bar=1 mm. Except **A**, data are represented as mean values ± SEM. P values were calculated using unpaired two-tailed student’s t test with Holm correction (**A**), unpaired two-tailed student’s t test (**C, F**), two-way ANOVA with Tukey multiple-comparison test (**E**) or one-way ANOVA with Tukey multiple-comparison test **(H)**. *, p<0.05; **, p< 0.01; ***, p< 0.001; ****, p< 0.0001; ns, non-significant.

To investigate whether the increased *Sca-1* expression within the tumor mass was due to an increased gene expression in all cancer cells or to the enrichment of a Sca-1^+^ population, we orthotopically injected 4T1 tumor cells and determined the frequency of different cell populations present in the primary tumor and lung metastases 30 days later by flow cytometry (Figure 1B). We observed that the frequency of both Sca-1^+^ and CD61^+^ populations increased in lung metastases compared to primary tumors (Figure 1C). In contrast, the CD24^+^, CD44^+^ and CD49f^+^ populations were not significantly altered.

The enrichment of the Sca-1^+^ population in lung metastases prompted us to ask whether Sca-1^+^ cells actively contribute to the metastasis. To this end, we isolated 4T1-Sca-1^+^ and 4T1-Sca-1^-^ cells by magnetic activated cell sorting (MACS) from parental 4T1 cells, which contains low frequency of Sca-1^+^ population (10-15%) (Supplemental Figure 1B, C), and examined their metastatic ability *in vivo*. In the orthotopic injection model, 4T1-Sca-1^+^ cells formed significantly more lung metastases than 4T1-Sca-1^-^ cells, while there was no significant difference in primary tumor growth (Figure 1D-F). Upon tail vein injection, 4T1-Sca-1^+^ cells displayed significantly greater lung colonization ability compared to 4T1-Sca-1^-^ cells and a non-significant increase compared to parental 4T1 cells (Figure 1G-I). In addition, 4T1-Sca-1^+^ cells showed significantly higher *in vitro* mammosphere forming efficiency than 4T1-Sca-1^-^ cells (Supplemental Figure 2A), while *in vitro* cell growth and cell motility were comparable (Supplemental Figure 2B-C).

These results suggest that 4T1-Sca-1^+^ and 4T1-Sca-1^-^ cells have similar tumorigenic potential, while 4T1-Sca-1^+^ cells have higher metastasis-initiating capacity.

### The Sca-1^+^ tumor cell population is plastic *in vitro* and *in vivo*

Growing evidence indicates that cancer cells possess plastic features, which can be modulated by both cell-intrinsic factors and microenvironmental cues (37, 38). To characterize the observed plasticity of 4T1-Sca-1^+^ cells, we first investigated isolated 4T1-Sca-1^+^ and 4T1-Sca-1^-^ cells *in vitro*. The abundance of Sca-1^+^ population in 4T1-Sca-1^+^ cells enriched by MACS sorting (> 75%) gradually decreased to 50% after 4 days of culture (Supplemental Figure 2D, upper panel), while the Sca-1^-^ cells (negatively enriched by MACS) regenerated a Sca-1^+^ population (from less that 1% to 19%) (Supplemental Figure 2E, lower panel). Consistently, after orthotopic injection of 4T1-Sca-1^+^ cells, the abundance of the Sca-1^+^ population in the derived tumors decreased from 75% to about 40% after 23 days of growth, while in tumors generated from 4T1-Sca-1^-^ cells it increased from less than 1% to 15% (Supplemental Figure 2E), similar to the frequency of Sca-1^+^ population in tumors derived from parental 4T1 cells (Figure 1C). In addition, when tumor cells derived from primary tumors and lung metastases of parental 4T1-injected mice were cultured *ex vivo*, the abundance of the Sca-1^+^ population significantly decreased from 20% to 4.5% and 60% to 19%, respectively (Supplemental Figure 2F).

From these observations we conclude that both Sca-1^+^ and Sca-1^-^ populations are highly plastic and this plasticity appears to be modulated *in vivo*.

### Tumor-educated Gr1^+^CD11b^+^ cells expand the metastatic Sca-1^+^ population

Immune cells in the TME are critical determinants of tumor cells functions and behaviors, including metastatic capacity (39). To collect evidence for a potential correlation between immune cells and the Sca-1^+^ population, we characterized the inflammatory cells infiltrating the orthotopic primary tumors. We observed a significant increase of the Gr1^high^CD11b^+^Ly6C^low^ population and a significant decrease of the Gr1^low^CD11b^+^Ly6C^high^ population in tumors derived from the 4T1-Sca-1^+^ cells, compared to tumors derived from the 4T1-Sca-1^-^ cells (Figure 2A). Gr1^high^CD11b^+^Ly6C^low^ cells are immature myeloid progenitors mobilized from the bone marrow by tumor-derived signals capable of establishing an immunosuppressive environment facilitating tumor progression and metastasis (40, 41). Gr1^+^CD11b^+^ cells form a homogenous population in the circulation (Supplemental Figure 3A), consistent with the literature (42), while in the TME they differentiated into two distinct subpopulations, Gr1^high^ and Gr1^low^. To examine the direct contribution of the tumor-educated Gr1^+^CD11b^+^ cells in promoting the enrichment of the Sca-1^+^ population, we isolated Gr1^+^ cells from primary tumors, bone marrow and spleen of 4T1 tumor-bearing BALB/c mice by MACS, and co-cultured them *in vitro* with parental 4T1 cells (Figure 2B). Gr1^+^ cells isolated from tumors (Tu-Gr1^+^CD11b^+^), but not from spleen (Spl-Gr1^+^CD11b^+^) or bone marrow (BM-Gr1^+^CD11b^+^), significantly induced the expansion of a Sca-1^+^ population in 4T1 cells (from 12.5% to 81%, p<0.001) in 48 hours (Figure 2C). To test whether the expansion of a Sca-1^+^ population from 4T1 cells required direct contact with Tu-Gr1^+^CD11b^+^ or was mediated by soluble factors, we compared the induction in two different co-culture setups, either in standard wells (cell contact) or in Transwells, where Tu-Gr1^+^CD11b^+^ and 4T1 cells were separated by a filter with 0.4 µm pores (Figure 2D). We did not observe any significant difference in the induction efficiency of Sca-1^+^ populations (measured by flow cytometry) between the two conditions and increasing the 4T1:Tu-Gr1^+^CD11b^+^ cell ratio from 1:1 to 1:3 did not further expand the Sca-1^+^ population (Figure 2E). Furthermore, conditioned medium from co-cultured Tu-Gr1^+^CD11b^+^ and 4T1 cells was also capable of subsequently expanding the Sca-1^+^ population from 4T1 cells alone (Supplemental Figure 3B). Interestingly, the Tu-Gr1^+^CD11b^+^ induced Sca-1^+^ population appeared more stable in time compared to the isolated 4T1-Sca-1^+^ cells (Supplemental Figure 3C and Supplemental Figure 2E). More importantly, when Gr1^+^CD11b^+^ cells-primed 4T1 cells were injected into the tail vein, Tu-Gr1^+^CD11b^+^ primed ones showed higher lung colonization capacity compared to Spl-Gr1^+^CD11b^+^ primed one (Figure 2F-H).

**Figure 2.**
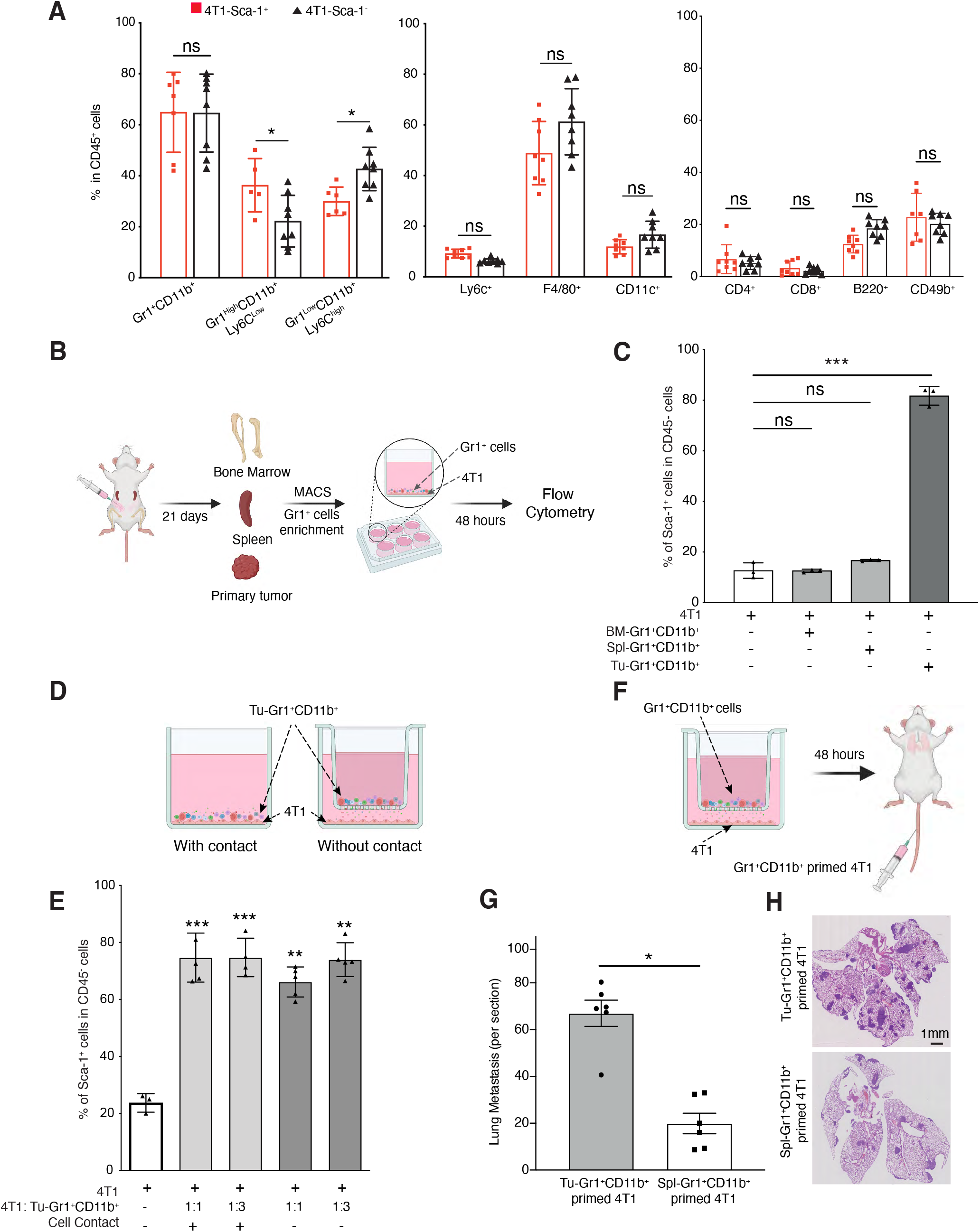
Sca-1 expression is modulated by the tumor microenvironment. **(A)** Frequency of different immune cell populations in primary tumors of mice orthotopically injected with 4T1-Sca-1+ and 4T1-Sca-1- 21 days post injection. Populations are determined in CD45 negative, viable cells (n=8 mice/Group). **(B-C)** Illustrative scheme **(B)** showing the experimental design for isolating Gr1+ cells from different sites of tumor-bearing mice. Twenty-one days after tumor implantation, Gr1+ cells were isolated from bone marrow (BM-Gr1+CD11b+), spleen (Spl-Gr1+CD11b+) or primary tumor (Tu-Gr1+CD11b+) and co-cultured for 48 hours with parental 4T1 cells in vitro. Sca-1 expression in tumor cells was examined by flow cytometry **(C)**. Co-cultures conditions are indicated in the bar graph (n=3/group). **(D)** Illustrative scheme of the experimental co-culture setup. **(E)** MACS-enriched Gr1+ cells were cocultured with 4T1 cells with or without Transwell inserts of 0.4 μm pore size. The 4T1 cells were seeded in the bottom well and Gr1+CD11b+ cells in the upper part of the insert. After 48 hours, 4T1 cells were examined for Sca1 expression by FACS. Co-cultures conditions are indicated in the bar graph. The ratio of tumor cells and Tu-Gr1+CD11b+ is varied from 1:1 to 1:3. Significant enrichment of Sca-1+ population were observed in all conditions. n= 3-5/group. **(F)** Illustrative scheme of the experimental metastasis setup. **(G-H)** Evaluation of the metastatic capacity of Gr1+CD11b+-educated 4T1 cells in vivo. The 4T1 tumor cells were primed with Tu-Gr1+CD11b+ or Spl-Gr1+CD11b+ in vitro without cell-cell contact for 48 hours and injected into the tail vein of mice. Lung metastases were quantified 10 days after injection **(G)**, and representative H&E staining images of lung sections are showed **(H)**. Scale bar=1mm, n=7 mice/group. Data are represented as mean values ± SEM from at least 3 independent experiments. P values were calculated using unpaired two-tailed student’s t test (**A, G**), one-way ANOVA with Dunnett’s multiple-comparison test (**C, E**). *, p<0.05; **, p< 0.01; ***, p< 0.001; ns, non-significant

These results imply that tumor-educated Gr1^+^CD11b^+^ cells induce the emergence of a highly metastatic Sca-1^+^ population through secreted factors.

### Tu-Gr1^+^CD11b^+^-induced and tumor-inherent Sca-1^+^ populations display distinct gene expression profiles

To unravel the molecular basis for the metastatic capacity of the inherent 4T1-Sca-1^+^ population and the Tu-Gr1^+^CD11b^+^ induced Sca-1^+^ population, we first performed transcriptomic profiling of 4T1-Sca-1^+^ and 4T1-Sca-1^-^ cells isolated from the parental 4T1 line. Pathway enrichment analysis showed that 4T1-Sca-1^+^ and 4T1-Sca-1^-^ cells expressed different genes associated with distinct signaling pathways (Figure 3A). The top 200 significantly upregulated and downregulated genes were extracted as Sca1 Positive and Sca1 Negative signatures, respectively (Supplemental Table 1). Next, we performed transcriptomic profiling of Tu-Gr1^+^CD11b^+^-primed 4T1, Spl-Gr1^+^CD11b^+^-primed 4T1 and parental 4T1 cells. Pathway enrichment analysis revealed that Tu-Gr1^+^CD11b^+^ and Spl-Gr1^+^CD11b^+^ priming induced distinct transcriptomic alterations in 4T1 cells (Figure 3B). To focus on the transcriptomic alternations related to the Sca-1^+^ population conversion, we compared Tu-Gr1^+^CD11b^+^-primed vs Spl-Gr1^+^CD11b^+^-primed cells. Interestingly, Tu-Gr1^+^CD11b^+^-primed 4T1 cells expressed both Sca-1 Positive and Sca-1 Negative signatures (Figure 3C), suggesting that Tu-Gr1^+^CD11b^+^ may twist tumor plasticity by converting the 4T1-Sca-1^-^ cells into 4T1-Sca-1^+^ cells, rather than expanding the pre-existing 4T1-Sca-1^+^ population.

**Figure 3.**
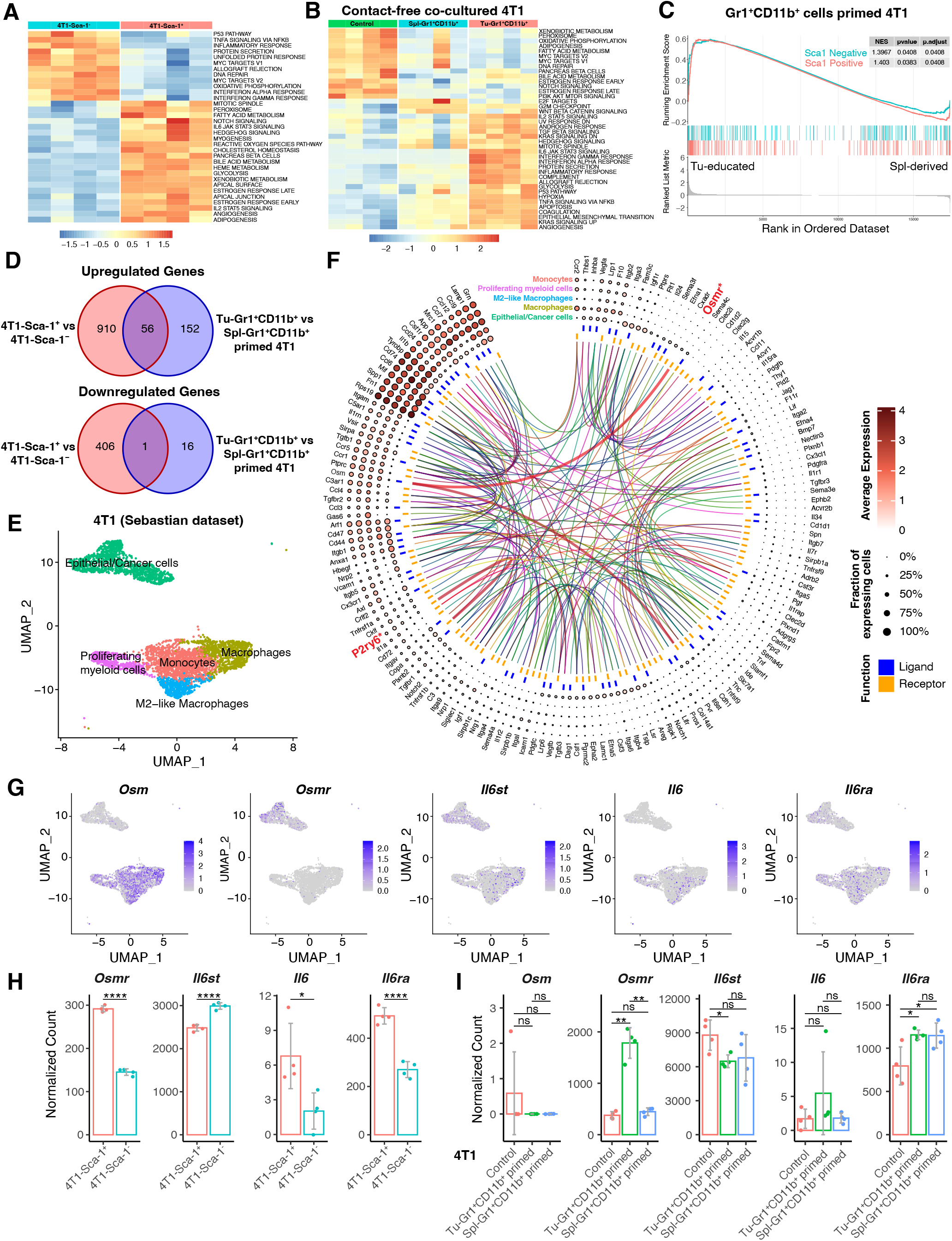
Transcriptomic analysis of Sca-1+ tumor cells. **(A)** Heatmap showing the signature score of the hallmark pathways analysis in 4T1-Sca-1+ and 4T1-Sca-1- population sorted from parental 4T1 cells. The colors code the expression levels relative to average levels as indicate at the bottom. **(B)** Heatmap showing the signature score of the hallmarks pathway analysis in parental 4T1 (4T1), Spl-Gr1+CD11b+ primed 4T1 and Tu-Gr1+CD11b+ primed 4T1 cells. The colors code the expression levels relative to average levels as indicate at the bottom. **(C)** Gene set enrichment analysis (GSEA) comparing the Tu-Gr1+CD11b+ and Spl-Gr1+CD11b+ primed 4T1 cells. GSEA shows positive correlations of both Sca1 Positive and Sca1 Negative signatures. NES, normalized enrichment score. **(D)** Venn diagrams showing that 56 upregulated genes, and 1 downregulated gene are shared between endogenous and Tu-Gr1+CD11b+ induced Sca-1+ population in 4T1 tumor cells. **(E)** UMAP plot showing clusters of cancer cells and myeloid cell populations in orthotopically growing 4T1-derived primary tumors extracted from the Sebastian dataset (see Materials and Methods for details). **(F)** Circos diagram showing the potential interactions between cancer cells and different myeloid cell populations determined by CellPhoneDB (see Materials and Methods for details) based on the Sebastian dataset. Only OSMR and P2RY6 are shared with the common 56 gene list showed in panel **D**. **(G)** UMAP plot showing the gene expression pattern of *Osm*, *Osmr*, *Il6st*, *Il6* and *Il6ra* in different cell populations in the Sebastian dataset. **(H-I)** mRNA expression of *Osm*, *Osmr*, *Il6st*, *Il6* and *Il6ra* based on RNAseq data used to generate the heatmaps shown in **A** & **B**, respectively. *Osm* expression is not detected in sorted 4T1 cells. Data are presented as mean of normalized count ± SD. P values: *, p<0.05; **, p< 0.01; ***, p< 0.001; ****, p< 0.0001; ns, non-significant (unpaired two-tailed student’s t test for **H**, and unpaired two-tailed student’s t test with Holm correction for **I**).

To test whether, despite the diverse transcriptional profiles across different Sca-1^+^ populations, there could be a common molecular mechanism underlying their induction and metastatic capacity, we compared the significantly differentially expressed genes between 4T1-Sca-1^+^ versus 4T1-Sca-1^-^ and Tu-Gr1^+^CD11b^+^ primed 4T1 versus Spl-Gr1^+^CD11b^+^ primed 4T1 cells. Strikingly, among a total of 1118 up- and 423 down-regulated genes found when comparing the two conditions, only 56 up- and one down- regulated genes were shared (Figure 3D and Supplemental Table 1). This observation was consistent with the notion that the Sca-1^+^ population in Tu-Gr1^+^CD11b^+^ primed 4T1 cells was different from the inherent 4T1-Sca-1^+^ cells. Nonetheless, the fact that the 4T1-Sca-1^+^ cells and Tu-Gr1^+^CD11b^+^ primed 4T1 cells possessed similar *in vivo* metastatic capacity suggested that among these common pathways some were relevant for 4T1 metastases formation. To this end, we analyzed the publicly available scRNA-seq dataset from 4T1 primary tumors of Sebastian *et al*. (43). In this dataset, several cell types, including cancer cells, epithelial cells, fibroblasts, distinct subpopulations of myeloid cells were identified (43). To determine significant ligand-receptor interactions from the scRNA-seq data, we performed cell-cell interaction analysis with CellPhoneDB (44) by focusing on the interactions between epithelial/cancer cells and myeloid cells (Figure 3E, F). The analysis identified 160 ligand-receptor interaction pairs (Supplemental Table 2). Among those pairs, OSM receptor (OSMR) and pyrimidinergic receptor P2Y6 (P2RY6) were the only ones present inside the 56 common genes shown in Figure 3D. However, P2RY6 interacts with COPA (Coatomer Complex Subunit Alpha), a membrane protein involved in membrane traffic between endoplasmic reticulum and Golgi (45) and, thus, unlikely to mediate cell-cell contact-independent induction of the Sca-1^+^ population. As the IL6-JAK-STAT3 signaling pathway was upregulated both in the 4T1-Sca-1^+^ population and Tu-Gr1^+^CD11b^+^-primed 4T1 cells (Figure 3A, B), we next examined the expression of *Osm*, *Osmr*, *Il6st*, *Il6*, and *Il6* receptor (*Il6ra*) in the Sebastian dataset (Figure 3G). The expression of *Il6* and *Osm* was restricted to myeloid cells, with *Osm* expression being more prominent, similar to a previous report (33). *Osmr* was predominantly expressed in tumor cells, while *Il6st* and *Il6ra* were homogenously expressed in all cell types.

We then explored their expression in the 4T1-Sca-1^+^ cells and Tu-Gr1^+^CD11b^+^ primed 4T1 cells. *Osm* and *Il6* expression were very low in all samples (normalized count number less than 7 on average) (Figure 3H-I), consistent with data in the Sebastian dataset (Figure 3G). *Osmr* and *Il6ra*, however, were highly expressed in 4T1-Sca-1^+^ cells compared with 4T1-Sca-1^-^ cells, while the expression of *Il6st* was abundant in both populations, though higher in 4T1-Sca-1^+^ cells (Figure 3H). On the other hand, only *Osmr* was significantly upregulated in Tu-Gr1^+^CD11b^+^ primed 4T1 compared with Spl-Gr1^+^CD11b^+^ primed 4T1 (Figure 3I).

Taken together, these results suggest that OSM/OMSR and IL6/IL6R signaling pathways may be involved in the Tu-Gr1^+^CD11b^+^-mediated expansion of 4T1-Sca-1^+^ cells.

### Tu-Gr1^+^CD11b^+^ cells promote Sca-1^-^ to Sca-1^+^ population conversion

The above results strongly implied that Tu-Gr1^+^CD11b^+^ cells convert Sca-1^-^ population into the Sca-1^+^ one. To further test this hypothesis, we compared the cell population dynamics in cultured cells and the orthotopic primary tumors by analyzing publicly available scRNA-seq datasets. By integrating scRNA-seq data from 3D cultured 4T1 cells (GSM4812003) (46) and tumor cells isolated from orthotopically fat pad-injected 4T1 primary tumor (PT) (GSM3502134) (47) (Figure 4A, B), we observed 5 clusters. Clusters 0, 1, 2, 4 were predominant in cultured tumor cells, while cluster 3 was predominant in primary tumors. Single-cell trajectories analysis confirmed that cluster 3 was at the end of the transformation process (Figure 4C, D). The population dynamics also showed that the fraction of cells in clusters 1, 2 and 4 decreased during the transformation, the one in cluster 0 only minimally increased, while the fraction in cluster 3 massively increased (Figure 4E). Importantly, very few cultured 4T1 cells expressed *Sca-1* while it was abundantly expressed in the majority of cells in the primary tumor (Supplemental Figure 4A, upper panel). Consistently, the fraction of cells expressing *Osmr* was higher in the primary tumor compared to cultured cells (Supplemental Figure 4A, lower panel). Similar observations were obtained when analyzing scRNA-seq data from the ER^+^ human breast cancer model MCF-7. After integrating data from cultured MCF-7 cells (GSM4681765) and tumor cells that were isolated from MCF-7 intraductal injected mammary gland (GSM5904917) (48), 6 clusters were identified (Figure 4F), with clusters 1 and 3 predominant in cultured MCF-7 cells, while clusters 2 and 4 were predominant in primary tumors (Figure 4G). Further analysis showed that clusters 2 and 4 expanded during *in vitro* to *in vivo* tumor cell transformation and represented nearly 50% of the *in vivo* primary tumor cells (Figure 4H-J). Although there is no human homolog of *Sca1* gene, *OSMR* expressing cells were increased upon tumor implantation, especially in clusters 2 and 1 (Supplemental Figure 4B).

**Figure 4.**
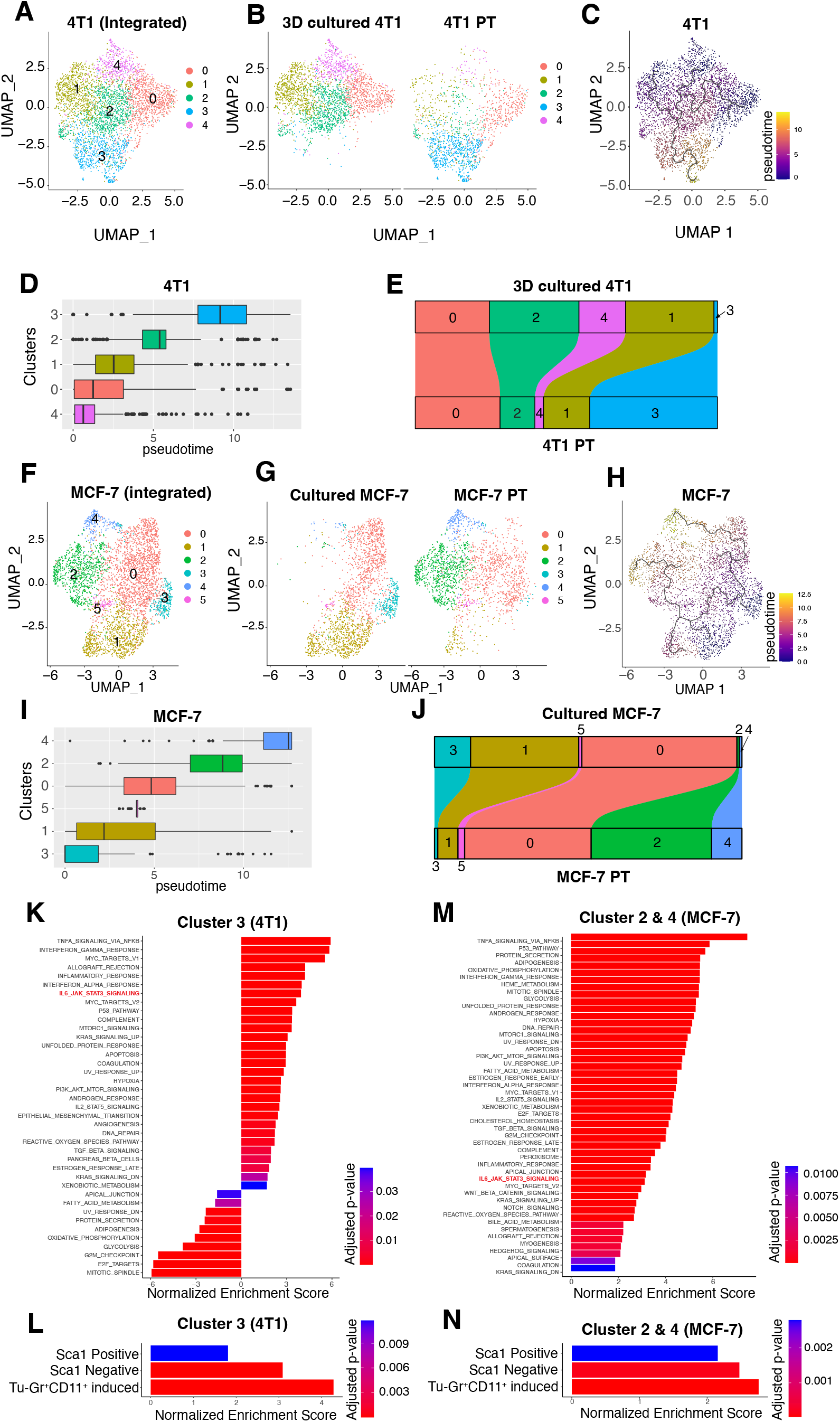
Transformation dynamics of tumor cell populations induced by the tumor microenvironment. **(A)** UMAP plots showing 4T1 clusters based on integrated scRNA-seq data from 4T1 cells in 3D culture or in primary tumor. **(B)** Distribution of specific clusters in 4T1 cells in 3D culture or in primary tumor (PT). **(C-D)** UMAP plot **(C)** and boxplot **(D)** showing the clusters in pseudo-time during the transformation of 4T1 cells from *ex vivo* culture to *in vivo*. **(E)** Sankey diagram showing the dynamic of each cluster during the transformation of 4T1 cells from *ex vivo* culture to in vivo. Cluster 3 was largely expanded *in vivo*. **(F)** UMAP plots showing MCF-7 clusters based on integrated scRNA-seq data from MCF-7 cells in culture or in primary tumor. **(G)** Distribution of the specific clusters in cultured MCF7 cells or in MCF7 primary tumors (PT). **(H-I)** UMAP plot **(H)** and boxplot **(I)** showing the clusters in pseudo-time during the transformation of MCF-7 cells from *ex vivo* culture to *in vivo.* **(J)** Sankey diagram showing the dynamic of each cluster during the transformation of MCF-7 cells from *ex vivo* culture to *in vivo*. Cluster 2 and 4 were largely expanded *in vivo*. **(K-L)** GESA analysis of Hallmark gene sets **(K)** and Sca1 Positive signature, Sca1 Negative Signature and Tu-Gr1+CD11b+ induced signature **(L)** of cluster 3 in 4T1 data. Only the signatures with adjusted p-value <0.05 were shown. **(M-N)** GESA analysis of Hallmark gene sets (M) and Sca1 Positive signature, Sca1 Negative Signature and Tu-Gr1+CD11b+ induced signature **(N)** of cells in cluster 2 or cluster 4 in MCF-7 data. Only the signatures with adjusted p-value <0.05 were shown. Analyses are based on publicly available data (4T1: GSM4812003 and GSM3502134; MCF-7: GSM4681765 and GSM5904917).

To further investigate the signals involved in this transformation, we performed GSAE analysis for cluster 3 in 4T1 cells and clusters 2 and 4 in MCF-7 cells, respectively (Figure 4K-N). By comparing the Hallmark gene signatures, IL6-JAK-STAT3 signature was significantly upregulated in both cell populations (Figure 4K, M). Interestingly, Sca1 Positive and Sca1 Negative signatures were both upregulated (Figure 4L, N), which is consistent with our *ex vivo* induction experiment (Figure 3C). To validate the involvement of Tu-Gr1^+^CD11b^+^ during the cell population transformation, we extracted the top 50 upregulated genes (Supplemental Table 3) identified by comparing the Tu-Gr1^+^CD11b^+^ with the Sp-Gr1^+^CD11b^+^-stimulated 4T1 cells as Tu-Gr1^+^CD11b^+^ induced signature. Both cell populations predominant in the primary tumor in both 4T1 and MCF-7 models, upregulated the Tu-Gr1^+^CD11b^+^ induced signature (Figure 4L, N).

These data, together with our *in vivo* observations (Figure 1A-C) and *ex vivo* coculture experiments (Figure 2B-E and Figure 3C), indicate that Tu-Gr1^+^CD11b^+^ convert the Sca-1^-^ population to Sca-1^+^ population, likely, via OSM/IL6 signaling pathway.

### OSM/IL6-JAK pathway mediates Tu-Gr1^+^CD11b^+^-induced Sca-1^+^ population enrichment

To experimentally interrogate the role of OSM/IL6 in modulating the Sca-1^+^ population, we first measured *Osm* and *Il6* mRNA expression in Spl-Gr1^+^CD11b^+^ and Tu-Gr1^+^CD11b^+^. Indeed, both *Osm* and *Il6* mRNA levels were significantly elevated in Tu-Gr1^+^CD11b^+^ (Figure 5A). To functionally validate the role of OSM/IL6 in the generation of Sca-1^+^ population, we treated 4T1 cells directly with recombinant OSM and IL6 proteins *in vitro* and measured the effect on the Sca-1^+^ population. After 2 days of treatment, both OSM and IL6 significantly increased the frequency of Sca-1^+^ cells from 16% to 38.8% and 26.5%, respectively (Figure 5B). Conversely, blocking OSM and IL6 activities using anti-OSM or -IL6 neutralizing antibodies significantly reduced the emergence of Sca-1^+^ population induced by Tu-Gr1^+^CD11b^+^ conditioned medium (Figure 5C). The combination of anti-OSM and -IL6 antibodies did not have additive effects suggesting that OSM and IL6 both contribute in promoting Sca-1^+^ population by sharing the same signaling cascades.

**Figure 5.**
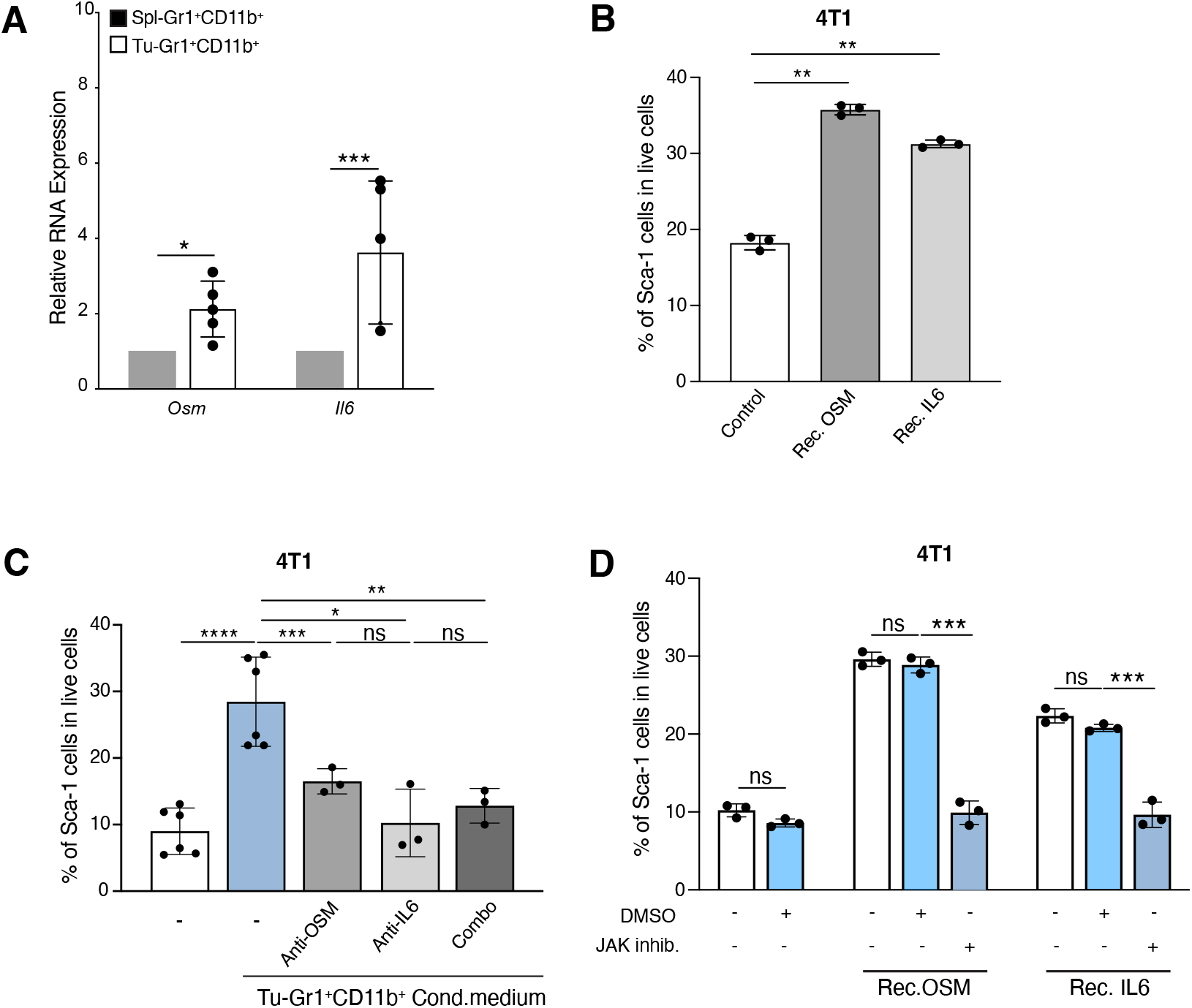
Sca-1+ population is modulated by the OSM/IL6-JAK pathway. (**A**) Quantitative PCR analysis of *Osm* and *Il6* mRNA expression in Tu-Gr1+CD11b+ and Spl- Gr1+CD11b+. Tu-Gr1+CD11b+ express significantly higher *Osm* and *Il6* levels compared with Spl-Gr1+CD11b+. n = 4-5/group. (**B**) Fraction of 4T1-Sca1+ cells upon exposure to recombinant Il6 or Osm protein (1 μg/ml for 48 hours) as determined by flow cytometry. Both cytokines induced the Sca-1+ population in cultured 4T1 tumor cells. n = 3/group. (**C**) Inhibition of OSM and IL6 from Tu-Gr1+CD11b+ conditioned medium with anti-OSM or anti-IL6 neutralizing antibody as indicated. Treatment with either antibody significantly suppressed the Sca-1+ population enrichment. n = 3-6/group. (**D**) Treatment with the JAK inhibitor Ruxolitinib (5 μM) of cultured 4T1 cells stimulated with recombinant IL6 or OSM protein (1 μg/ml, 48 hours exposure) as indicated. Ruxolitinib inhibited Sca-1+ population enrichment induced by recombinant IL6 or OSM protein. n = 3/group. Data are represented as mean ± SEM from 3 independent experiments. *, p<0.05; **, p< 0.01; ***, p< 0.001; ****, p< 0.0001; ns, non-significant (unpaired two-tailed student’s t test for **A**, and one-way ANOVA with Dunnett’s multiple-comparison test for **B** and **D**, and one-way ANOVA with Tukey multiple-comparison test for **C**).

OSMR and IL6R signal by activating the intracellular Janus tyrosine kinase (JAK) (49). To explore the involvement of the JAK pathway in the emergence of the Sca-1^+^ population, we treated 4T1 cells with the JAK inhibitor (Ruxolitinib) during exposure to recombinant OSM and IL6. Ruxolitinib treatment prevented the emergence of the Sca-1^+^ population in response to recombinant OSM and IL6 (Figure 5D).

From these results, we conclude that OSM/IL6-JAK pathway mediates Tu-Gr1^+^CD11b^+^ -induced Sca-1^+^ population enrichment.

### Tu-Gr1^+^CD11b^+^-induced Sca-1^+^ population and 4T1-inherent Sca-1^+^ population have distinct CSC and EMT gene expression profiles

OSM/IL6-JAK signaling has been reported to support tumor progression by promoting a CSC phenotype and epithelial-mesenchymal plasticity (25, 36, 50). To further characterize CSC and EMT features in 4T1-Sca-1^+^ cells and Tu-Gr1^+^CD11b^+^ induced Sca-1^+^ population, we interrogated our RNAseq data set for the expression of 17 stem cell and EMT markers (Supplemental Figure 5). The stem cell markers *Oct4* (*Pou5f1*), *Sox2* and *Nanog* were undetectable or very low in all or some samples. 4T1-Sca-1^+^ cells had higher expression of *Aldh1a1*, *Aldh3a1* and *Podxl* but lower expression of *Klf4* and *Sox9* compared to 4T1-Sca-1^-^ cells. There was no difference in the expression of *Abcg2* and *Has2*. Tu-Gr1^+^CD11b^+^ primed 4T1 cells had higher expression of *Klf4* and *Has2*, lower expression of *Aldh1a1*, *Aldh3a1* and *Sox9*, and similar expression of *Podxl* when compared with Spl-Gr1^+^CD11b^+^ primed 4T1. Among them, only *Has2* expression was specifically elevated in Tu-Gr1^+^CD11b^+^ primed 4T1 compared with control 4T1 and Spl-Gr1^+^CD11b^+^-primed 4T1 (Supplemental Figure 5A). On the other hand, 4T1-Sca-1^+^ cells had lower expression of *Cdh1* and higher expression of *Snail1*, *Twist1*, *Vim* and *Foxc1* which support an EMT status, although *Zeb1* expression was reduced (Supplemental Figure 5B). Globally, the expression of most of the EMT genes were similar between Tu-Gr1^+^CD11b^+^ and Spl-Gr1^+^CD11b^+^ primed 4T1 cells, except for *Snail2* and *Vim*, whose expression was suppressed in Tu-Gr1^+^CD11b^+^ primed 4T1 (Supplemental Figure 5B).

Taken together, these results indicate that the Tu-Gr1^+^CD11b^+^-induced Sca-1^+^ population and inherent Sca-1^+^ population have different CSC and EMT transcriptional profiles, reinforcing the notion that, Tu-Gr1^+^CD11b^+^-induced Sca-1^+^ population are not just enriched tumor-inherent Sca-1^+^ population

### Chemotherapy enriches a Sca-1^+^ population with CSC features

CSC and cancer cell plasticity contribute to drug resistance in various tumor types, including breast cancer (51–55). The above results, including the significantly elevated expression of *Aldh3a1* (a marker of drug resistance) in the 4T1-Sca-1^+^ population prompted us to investigate the resistance of this population to chemotherapy. To this end, we treated 4T1 cells for 48 hours *in vitro* with methotrexate (MTX) and doxorubicin (Dox), two widely used chemotherapy drugs, including in breast cancer. The 48 hours treatment of either drug increased the frequency of Sca-1^+^ population in 4T1 cells (Figure 6A). Next, we mimicked a clinically relevant situation of cancer cells escaping chemotherapy by exposing 4T1 cells to 28 nM MTX, a concentration slightly higher than the IC_50_ concentration of the drug, for up to 3 weeks and recovered the surviving cells by switching to normal medium (Figure 6B). The selected cell line, named MR13, was highly enriched in Sca-1^+^ cells (>60%) (Figure 6C). Compared to parental 4T1 tumor cells, MR13 cells exhibited a higher mammosphere forming efficiency (Figure 6D), lower *in vitro* proliferative capacity (Figure 6E) and increased cell mobility (Figure 6F), which were consistent with CSC-like properties. When tested in a 48-hours cytotoxicity assay, MR13 cells were more resistant against MTX compared to parental 4T1 (Supplemental Figure 6A).

**Figure 6.**
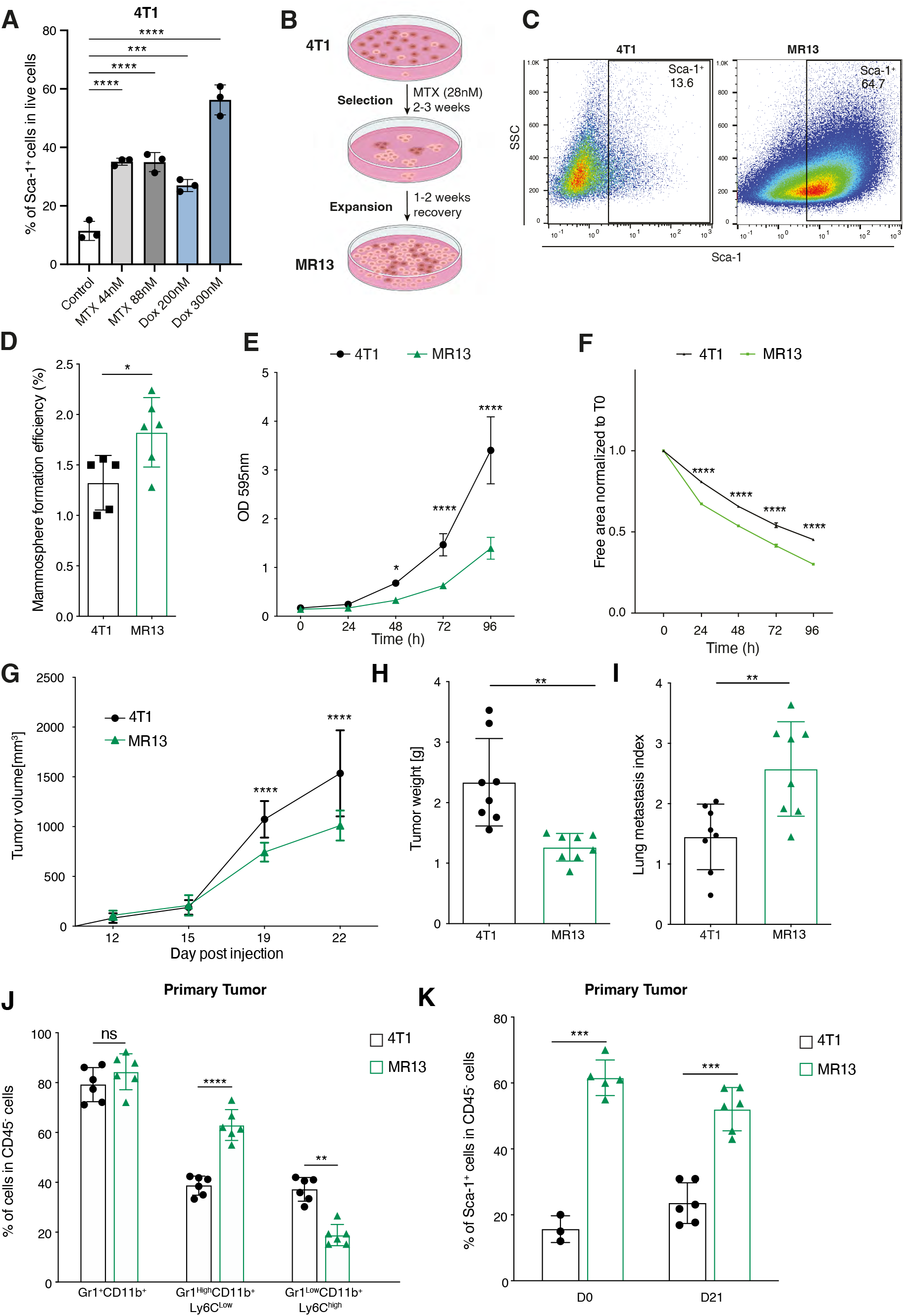
Long term chemotherapy treatment of 4T1 cells induces a stable Sca-1+ population (MR13) with higher metastatic capacity and CSC features. **(A)** Fraction of 4T1-Sca1+ cells upon short-term (48 hours) Methotrexate (44 and 88 nM) and Doxorubicin (200 and 300 nM) treatments. All treatments induced enrichment of Sca-1+ population. n = 3/group. **(B)** Illustrative scheme of the experimental design to obtain chemotherapy resistant MR13 cells from 4T1. **(C)** Dot plots representing Sca-1 expression vs SSC determined by flow cytometry in MR13 chemotherapy resistant cells vs parental 4T1 tumor cells in vitro. **(D)** Quantification of the mammosphere forming efficiency of 4T1 and MR13 tumor cells. n = 5-6/group. **(E)** Cell proliferation curve of 4T1 and MR13 tumor cells in vitro determined by crystal violet assay. The results are presented as mean of optical density (OD). n = 8/group. **(F)** Cell motility of 4T1 and MR13 tumor cells determined by a scratch wound healing assay. n=5-6/group. Results are presented as cell-free area relative to the initial wound area from 3 independent experiments. **(G)** Growth curves of primary tumors in BALB/c mice orthotopically injected with 4T1 and MR13 tumor cells (n=10-11/group). **(H)** Tumor weight of 4T1 and MR13 tumors recovered from BALB/c mice at day 22 post injection (n=8-9/group). **(I)** Lung metastasis index 23 days post injection. The number of metastatic nodules is determined by H&E staining and normalized based on the primary tumor weight (n=8-9/group). **(J)** Frequency of different CD11b+ myeloid cells subpopulations in primary tumors from MR13 and 4T1 injected mice determined by flow cytometry 21-days post injection (n= 6). Subpopulations are determined in CD45 positive, viable cell population. **(K)** Percentage of Sca-1+ tumor cells at time of injections (D0) of 4T1 and MR13 cells and in primary tumors recovered at day 21 (D21). Sca-1 expression is determined in CD45 negative, viable cell population. n = 3-6/group. Data are represented as mean ± SEM from at least 3 independent experiments. *, p<0.05; **, p< 0.01; ***, p< 0.001; ****, p< 0.0001; ns, non-significant (one-way ANOVA with Dunnett’s multiple-comparison test for **A**, unpaired two-tailed student’s t test for **D**, **H-K**, and two-way ANOVA with Tukey’s multiple comparison test for **E-G**).

### MR13-derived tumors are highly metastatic and rich in Gr1^high^CD11b^+^ Ly6C^low^ cells

To characterize the *in vivo* behavior of MR13 cells, we orthotopically implanted them into BALB/c mice and monitored the progression. MR13 cells formed smaller primary tumors relative to parental 4T1 cells, that were more metastatic to the lung (Figure 6G-I) and enriched in Gr1^high^CD11b^+^ Ly6C^low^ cells compared to 4T1 tumors (Figure 6J), similarly to 4T1-Sca-1^+^-derived tumors (Figure 2A). Strikingly, we observed metastases in the heart (Supplemental Figure 6B), which we never observed with the parental 4T1 cells. MR13 cells retained a large fraction of the Sca-1^+^ population *in vitro*, even when cultured in the absence of MTX (Figure 6C) and upon *in vivo* expansion (Figure 6K). Such stability of Sca-1 population contrasted with 4T1-Sca-1^+^ cells and Tu-Gr1^+^CD11b^+^-primed 4T1 cells (Supplemental Figure 2E, G) that, upon isolation or induction reverted to a Sca-1^-^ phenotype, suggesting that MR13 line was capable of self-sustain its own Sca-1^+^ population.

Taken together, chemotherapy-selected MR13 cells share some similar *in vivo* characteristics as 4T1-Sca-1^+^ cells, while they are capable of self-sustaining high Sca-1^+^ abundancy both *in vitro* and *in vivo*.

### IL6/IL6R-JAK autocrine signaling maintains Sca-1 positivity and metastatic capacity in MR13 cells

To better understand the chemotherapy-induced alterations in MR13 cells, we performed transcriptomic analyses comparing MR13 and parental 4T1 cells. Pathway enrichment analysis showed that the IL6-JAK-STAT3 signature was also elevated in MR13 cells (Figure 7A). Importantly, MR13 gene expression significantly positively correlated with the Sca1 Positive signature. At the same time, it negatively correlated with the Sca1 Negative signature (Figure 7B). This observation suggested that chemotherapy enriched for the inherent 4T1-Sca-1^+^ population, rather than converting plastic Sca-1^-^ cells, as observed upon Tu-Gr1^+^CD11b^+^-stimulation. In addition, *Osmr*, *Il6*, *Il6ra* and *Il6st* were all overexpressed in MR13 cells compared to parental 4T1 cells, while the *Osm* expression was not altered (Figure 7C). To functionally validate the IL6-JAK signaling pathway in Sca-1^+^ population maintenance, we treated MR13 cells with Ruxolitinib *in vitro* for 48 hours. The treatment significantly decreased the fraction of Sca-1^+^ cells (Figure 7D). Importantly, *in vitro* treatment of MR13 cells with Ruxolitinib for 3 days nearly completely abolished their lung metastatic capacity upon tail vein injection (Figure 7E-F).

**Figure 7.**
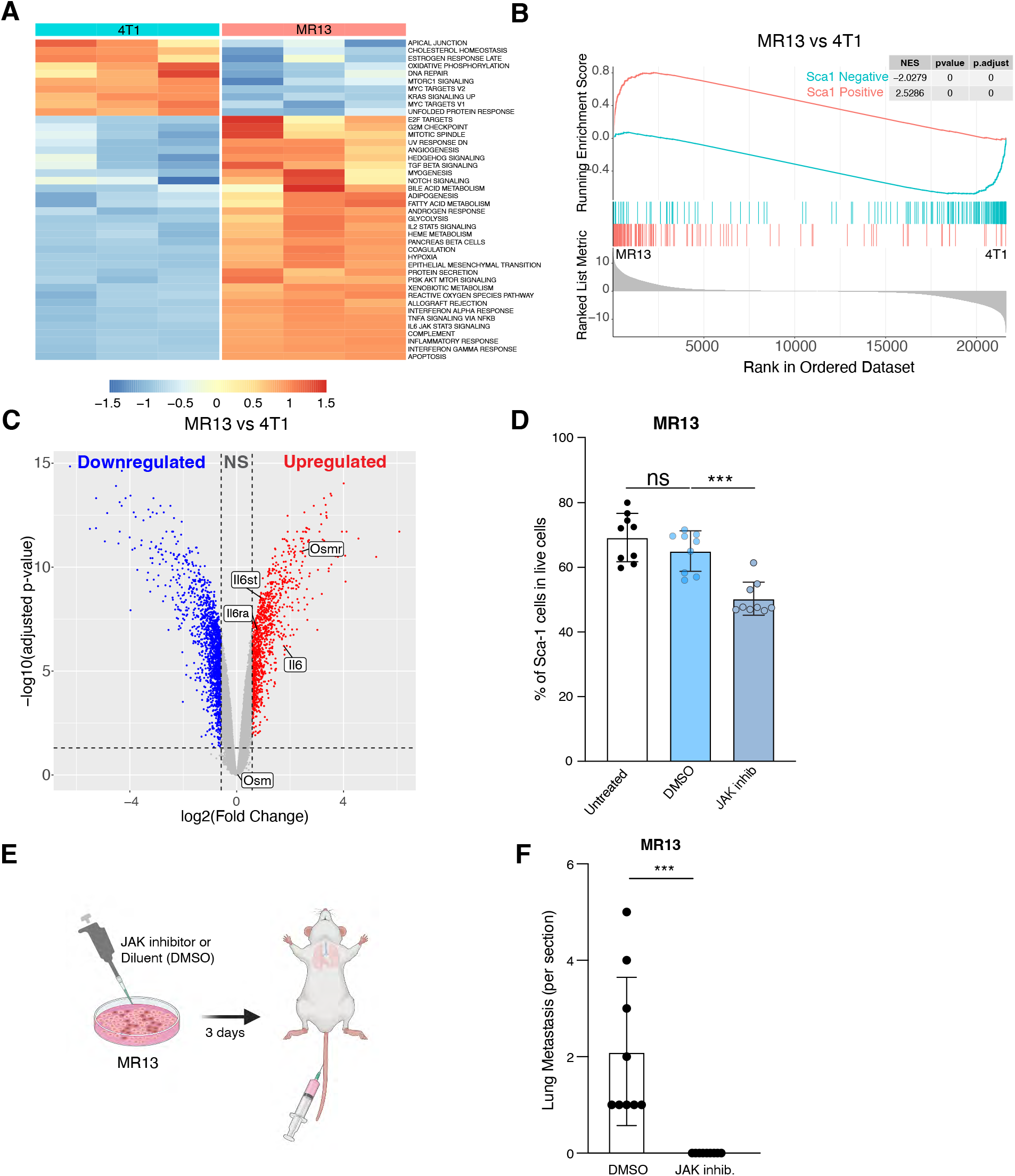
IL6-JAK pathway promotes Sca-1+ persistence and metastatic capacity in chemotherapy resistant MR13 tumor cells. **(A-B)** Gene expression analysis of parental 4T1 and chemotherapy resistant MR13 cells. Heat map represents the signature score of the hallmark pathways analysis. Results from 3 biological replicates are shown **(A)**. GSEA results showing that MR13 cells are positively enriched with for the Sca1 Positive signature and negatively with for the Sca1 Negative signature **(B)**. **(C)** Volcano plot showing the differential expression of *Osm*, *Osmr*, *Il6st*, *Il6* and *Il6ra* mRNA in MR13 vs 4T1 tumor cells. **(D)** Fraction of Sca-1+ population in MR13 tumor cells treated for 48 hours with Ruxolitinib (5 μM) relative to vehicle control (DMSO) treatment. **(E)** Illustrative scheme of the experimental design for testing the effect of Ruxolitinib on MR13 metastatic capacity shown in F. MR13 tumor cells were treated with Ruxolitinib or DMSO in vitro for 72 hours and then injected into the mice tail vein. Lungs were examined for metastasis 10 days after tumor cell injection. **(F)** Number of metastatic nodules in the lungs from mice injected with MR13 treated in vitro with Ruxolitinib or DMSO and indicated (n=9/group). Data are represented as mean values ± SEM from 3 independent experiments. P values: ns, non-significant, ***< 0.001 (one-way ANOVA with Dunnett’s multiple-comparison test for **D**, and unpaired two-tailed student’s t test for **F**).

Altogether, these data suggest that MR13 cells sustain the metastatic Sca-1^+^ population through cell-autonomous activation of the IL6-JAK signaling pathway.

### Tu-Gr1^+^CD11b^+^ invoked tumor cell signature predicts shorter overall and relapse-free survival in breast cancer patients

To evaluate the clinical relevance of the crosstalk between Tu-Gr1^+^CD11b^+^ and tumor cells, we tested whether the Tu-Gr1^+^CD11b^+^-induced 4T1 signature could predict cancer progression in patients. To this end, we interrogated the METABRIC dataset (56) with the Tu-Gr1^+^CD11b^+^-induced signature. Thirty-two human orthologue genes (Supplemental Table 3) in the murine 50 genes signature were present in the METABRIC dataset. Patients with the higher expression level of the signature had shorter overall survival (OS; p = 0.0056) and relapse-free survival (RFS; p = 0.032) (Figure 8A, B). Notably, OSM expression positively correlated with the signature in all patients (Supplemental Figure 7A), suggesting OSM do contribute to altered signature expression in patients. Of the thirty-two genes, five (*MX1, IRF7, OAS1, CMPK2, ISG15*) were commonly discriminant for a shorter OS and RFS (Figure 8C, D), and taken together, they further enhanced the predictive power (OS: p=0.00055; RFS: p=0.00069) (Figure 8E, F).

**Figure 8.**
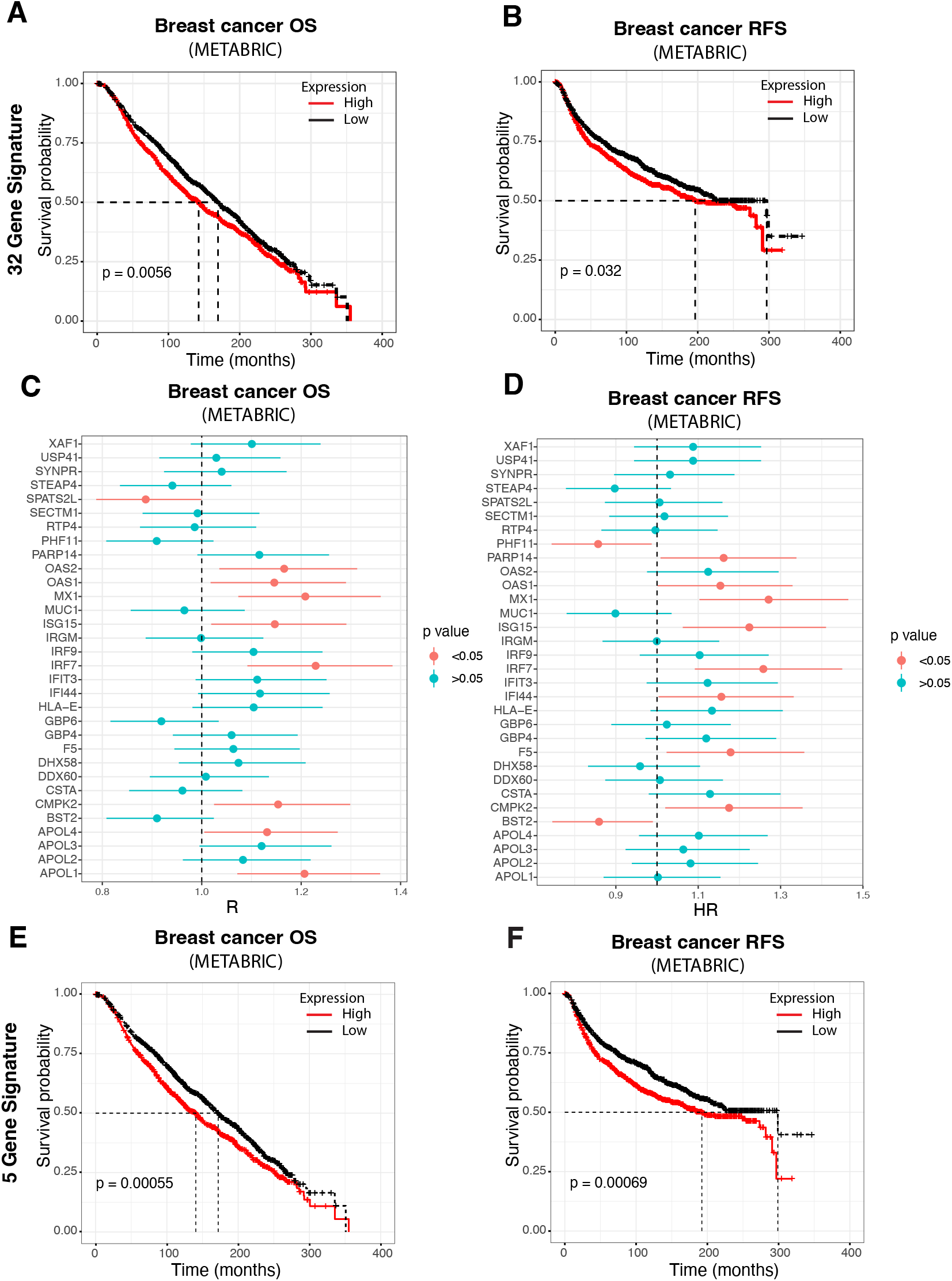
Tu-Gr1^+^CD11b^+^-induced tumor cell signature predicts worse outcome in breast cancer patients. (**A-B**) Kaplan-Meier curves showing overall survival (OS) (**A**) or relapse-free survival (RFS) (**B**) for breast cancer patients according to high or low expression of an orthologue 32 gene signature, based on the Tu-Gr1^+^CD11b^+^-induced 4T1 cell signature, in the METABRIC datasets. (**C-D**) Forest plots showing the Cox proportional hazard regression (HR) for OS (**C**) and RFS (**D**) of the individual 32 orthologues of the Tu-Gr1^+^CD11b^+^-induced signature, based on gene expression in tumor samples from METABRIC dataset. (**E-F**) Kaplan-Meier curves showing OS (**E**) and RFS (**F**) according to the reduced 5 orthologue gene signature expression in the METABRIC datasets. The p value for **A**, **B**, **E**, **F** was calculated using the log-rank test and high and low expression levels were stratified by median values.

These data indicate that a small set of genes issued from the murine tumor cell signature invoked by Tu-Gr1^+^CD11b^+^ cells can predict a shorter OS and RFS in breast cancer patients, thereby reinforcing the clinical significance of the proposed model.

## Discussion

Metastatic disease and therapy resistance are the leading causes of breast cancer mortality, calling for novel approaches to effectively prevent and cure for metastasis and therapy resistance. This is particularly relevant for TNBC, where in spite of the recent encouraging results with targeted therapies, such as PARP inhibitors for tumors with germline BRCA mutations, or checkpoint inhibitors for PD-L1^+^ tumors, management of metastatic disease remains challenging (57–60). It has been proposed that CSCs present in the primary tumor are responsible for tumor persistence, metastasis, and therapy resistance (52, 61). Enrichment and differentiation of CSC contribute to tumor heterogeneity (17, 19–22). Importantly, CSC features can be intrinsic or plastic and be modulated by cues from the TME (62–64).

Here, we have interrogated the contribution of the Sca-1^+^ population to breast cancer metastasis and its modulation by the TME. We reported that tumor-educated Gr1^+^CD11b^+^ cells (Tu-Gr1^+^CD11b^+^) instigate cancer metastasis by twisting cancer cell plasticity and enriching for a Sca-1^+^ population with enhanced metastatic capacity. We identified OSM/IL6-JAK as a paracrine communication axis between Tu-Gr1^+^CD11b^+^ and breast cancer cells and as an autocrine loop in chemotherapy-resistant tumor cells, promoting tumor heterogeneity, CSC features and metastatic capacity. Importantly, breast cancer patients expressing high levels of the human orthologues of the gene expression signatures invoked by Tu-Gr1^+^CD11b^+^ have significantly shorter OS and RFS, reinforcing the clinical significance of our findings. While some of these elements have been reported individually before, our results extend these observations by providing an integrative view of paracrine (Tu-Gr1^+^CD11b^+^-induced) and autocrine (chemotherapy-induced) communication in regulating tumor heterogeneity, cancer cell plasticity, metastasis, and resistance to chemotherapy.

A main observation stemming from this study is that Sca-1^+^ population exists under three different conditions: as inherent 4T1-Sca-1^+^ population, upon exposure to Tu-Gr1^+^CD11b^+^ cells and upon chemotherapy treatment (MR13). All these populations had higher metastatic abilities compared to their counterpart controls. GSEA analysis suggested that the Tu-Gr1^+^CD11b^+^-induced Sca-1^+^ population was likely to convert from the Sca-1^-^ population (Figure 3C and Figure 4), while the Sca-1^+^ population surviving chemotherapy (MR13) appeared to be enriched from the inherent Sca-1^+^ population (Figure 7B). In addition, the different gene expression signature of Tu-Gr1^+^CD11b^+^-induced Sca-1^+^ population relative to the inherent Sca-1^+^ cells suggest a remarkable functional plasticity of these cells. This plasticity is further supported by comparing single cell gene expression of murine 4T1 and human MCF-7 breast cancer cells, using publicly available scRNA-seq datasets, before and after *in vivo* growth (Figure 4). *In vivo*, tumor cells undergo a transformation enriching for Sca-1 (in the mouse), OSMR expressing cells, Sca1 Positive, Sca1 Negative and Tu-Gr1^+^CD11b^+^-induced signatures, and IL6-JAK signaling pathway from precursor cells (Sca-1^-^ in the mouse) (Figure 4). Gong *et al.* have reported that sorted Sca-1^-^ 4T1 cells could be transiently transformed into a Sca-1^+^ population by radiotherapy (65). An analogous observation was reported in colorectal cancer, where selective ablation of LGR5^+^ CSCs in organoids leads to initial tumor regression, followed by regrowth driven by LGR5^+^ CSCs reemerging from the LGR5^-^ population (66). Taken together, by demonstrating that tumor recruited and educated Gr1^+^CD11b^+^ cells contribute to such plasticity by inducing the conversion of low metastatic Sca-1-population into a highly metastatic Sca-1^+^ population, our observations further consolidate the notion that cancer consists of a heterogenous and plastic tumor mass, including highly metastatic cell populations. Dedicated time course scRNA-seq analyses together with lineage tracing experiments may help to further characterize the detailed origin, development, fate, and function of these Sca-1^+^ populations during cancer progression.

Recruitment and accumulation of Gr1^+^CD11b^+^ cells in the TME, particularly through the chemokines CCL2, CXCL1 and CXCL2, or IL-33, is considered a critical step for their contribution to tumor progression and metastasis (67–70). Consistent with these observations, tumors derived from sorted 4T1-Sca-1^+^ cells, or MR13 cells that are intrinsically enriched for Sca-1^+^ cells, have a higher content of Gr1^high^CD11b^+^Ly6C^low^ cells compared to tumors derived from 4T1-Sca-1^-^ or parental 4T1 cells, respectively (Figure 2A and 5K). Beyond recruitment, tumor-mediated education of Gr1^+^CD11b^+^ cells, appears to be necessary to gain higher metastatic activity: Only Gr1^+^CD11b^+^ cells recovered from primary tumors (but not from spleen or bone marrow) induced Sca-1 positivity and enhanced metastatic ability in 4T1 cells (Figure 2C, E). The role of Gr1^+^CD11b^+^ cells in promoting metastasis has been mainly attributed to promotion of angiogenesis, EMT and immunosuppression (17, 68). Peng *et al.*, reported the ability of those cells to endow CSC-like features to breast cancer cells but their metastatic capacity was not interrogated (17). Our observations suggest a self-sustaining positive feedback mechanism between highly metastatic cancer cells (Sca-1^+^ population) and Gr1^+^CD11b^+^ cells: inherent Sca1^+^ CSC-like, metastatic, cells promote recruitment and local education of Gr1^+^CD11b^+^ cells, which in turn promote tumor heterogeneity, cancer cell plasticity and metastatic capacity by converting low-metastatic Sca1^-^ cells into additional high-metastatic Sca1^+^ cells (Figure 9). Recently, it was reported that neutrophils escorting blood circulating tumor cells (CTCs) expands the metastatic potential of CTCs (71). While this effect was attributed to the promotion of cell cycle progression of CTCs through direct contact with the neutrophils, in light of our findings, one may also consider the possibility that clustered neutrophils may also promote the expansion of a CSCs-like phenotype with higher metastatic capacity. Interestingly, OSM was reported to be expressed by neutrophils cocultured with breast cancer cells (36) and to promote phenotypic changes associated with mesenchymal and stem cell-like differentiation in breast cancer (36, 72). Together with our observation, these findings further reinforce the notion that boosting the *Osm* expression in Gr1^+^CD11b^+^ cells is part of their educating program prompted by the tumor. One outstanding question raised by these observations is by which mechanisms and pathways cancer cells, in particular Sca-1^+^ ones, educate Gr1^+^CD11b^+^ cells to acquire cancer plasticity-promoting activity.

**Figure 9.**
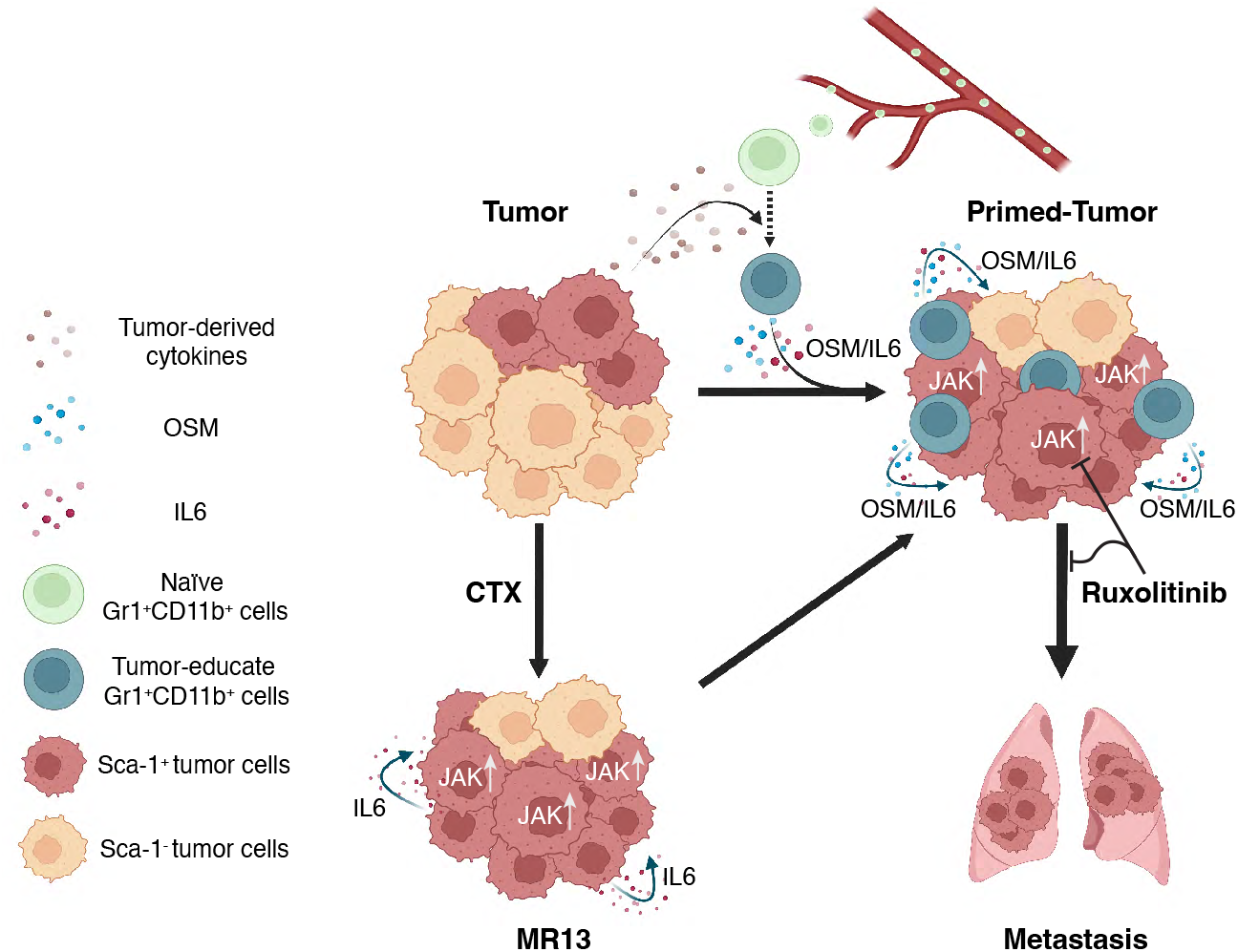
Illustrative scheme of the proposed model for cancer cell plasticity modulated by OSM/IL6 during tumor progression and chemotherapy. Parental tumor cells maintain a small portion of highly metastatic Sca-1^+^ population. During tumor progression, naïve Gr1^+^CD11b^+^ are recruited to the TME and educated into Tu-Gr1^+^CD11b^+^ by tumor-derived factors. In turn, Tu-Gr1^+^CD11b^+^ secrete OSM and IL6 to convert Sca-1^-^ population into a highly metastatic Sca-1^+^ population. Chemotherapy (CTX) enriched for Sca-1^+^ population due to its intrinsic resistance against cytotoxic treatment. Resistant cells express IL6 to maintain the high portion of Sca-1^+^ population with high metastatic ability. JAK inhibitor Ruxolitinib suppresses the conversion to Sca-1^+^ population and metastasis. Inhibition of OSM, IL6 and the activated downstream kinase JAK are candidate therapeutic targets to impinge on metastatic breast cancer progression during natural evolution and following therapy-resistance.

Besides directly activating tumor cells, OSM has also been shown to remodel macrophages and fibroblasts of the TME (33, 36). Araujo *et al.* recently reported that OSM derived from tumor-infiltrating myeloid cells reprogram fibroblasts to secrete VEGF and the chemokines CXCL1 and CXCL16, resulting in enhanced myeloid cell recruitment and breast cancer progression (33). Here we extend these observations by demonstrating that tumor-educated Gr1^+^CD11b^+^-derived OSM/IL6 twist cancer cell plasticity by promoting a rapid but reversible conversion of Sca-1^-^ cells into more metastatic Sca-1^+^ cells (Figure 5B). We further broaden the implications of the OSM/IL6-JAK axis, by demonstrating that MR13 cells that have escaped chemotherapy hijacked this paracrine mechanism in a cell autonomous manner by elevating the *Il6* expression (Figure 9). Significantly, a short *in vitro* treatment with Ruxolitinib effectively abrogated their metastatic capacity (Figure 7E, F).

While OSM/OSMR is the only interaction pair identified by the cell-cell interaction analysis (Figure 3F, G), it is possible that other molecular mechanisms may also play roles in modulating Sca-1^+^ cell plasticity and metastasis. One candidate is the IL6/IL6R communication axis, which was less prominent in the cell-cell communication analysis (Figure 3F, G), but highly expressed in our experimental data in Tu-Gr1^+^CD11b^+^ compared with Spl-Gr1^+^CD11b^+^ (Figure 5A) and in chemotherapy-resistant MR13 cells compared with 4T1 (Figure 7C). Moreover, the neutralization of IL6 also suppressed the Tu-Gr1^+^CD11b^+^ conditioned medium induced Sca-1^+^ cell enrichment, much alike OSM inhibition (Figure 5C).

Importantly, we demonstrated that a human orthologue signature of the 4T1 gene expression signature invoked by Tu-Gr1^+^CD11b^+^ can predict a significantly shorter OS and RFS in breast cancer patients (Figure 8). This finding strengthens the clinical significance of the observed crosstalk between Tu-Gr1^+^CD11b^+^ and tumor cells. Strikingly, the five genes that significantly contribute to the discriminatory power of the signature are genes related to native or viral immunity or regulated by interferon. While expression of interferons and interferon response genes in breast cancer has been mainly associated with tumor suppression and improved survival (73), there is evidence also correlating interferon responses with tumor promotion, therapy resistance and reduced survival (74). As JAKs/STATS are activated by both IFN and OSM/IL6 receptors, it is conceivable that OSM/IL6 only activates a subset of the IFN-induced genes with tumor-promoting activity, as the case for Mx1 (75). Consistent with our findings, JAK/STAT signaling has recently been shown to initiate the lineage plasticity in prostate cancer as well as to promote lineage plasticity-driven targeted therapy resistance in a stem-like subpopulation of prostate cancer (76, 77). On the other hand, Aouad *et al.* showed that epithelial-mesenchymal plasticity is essential for the generation of a dormant cell state of ER^+^ breast cancer during progression, and the activation of IL6-JAK-STAT signaling triggers tumor cell awakening and recurrence (48).

One crucial question raised by these results is whether the OSM/IL6-JAK pathway is a potential actionable clinical target to impinge on metastatic progression and therapy resistance. In particular, the observation that a short *in vitro* treatment of the highly metastatic MR13 cells profoundly suppressed their metastatic capacity *in vivo* (Figure 7E, F), suggests potential long-lasting effects consistent with an adjuvant effect. JAK inhibitors are being tested in clinical trials in breast cancer. A phase I study combining Ruxolitinib with paclitaxel in HER2-negative metastatic breast cancer showed good tolerability and evidence of activity (78). A phase I/II trial of Ruxolitinib in combination with trastuzumab in metastatic HER2 positive breast cancer and a phase II study combining Ruxolitinib with capecitabine in advanced HER2^-^ breast cancer, however, did not improve progression-free survival (79, 80). The absence of benefits in these studies in advanced breast cancer and our reported mechanistic observations on metastatic progression, raise the question of whether the JAK inhibitors should be considered in adjuvant setting in high-risk patients, to prevent progression to metastases, rather than treating patients already bearing metastases.

In conclusion, we reported here that a subpopulation of tumor cells within the tumor mass educates requited Gr1^+^CD11b^+^ cells to convert a low metastatic subpopulation into highly metastatic one, through the OSM/IL6-JAK signaling axis (Figure 9). The clinical relevance of this observation is supported by human transcriptomic data. This process is hijacked by tumor cells that survived chemotherapy and evolved toward a highly metastatic phenotype via cell autonomous IL6-JAK signaling. Importantly, a short *in vitro* treatment with a clinically approved JAK inhibitor, Ruxolitinib, suppresses their metastatic capacity these cells *in vivo.* These results should stimulate considering testing JAK inhibitors in the adjuvant setting in TNBC breast cancer patients at high-risk for metastatic progression.

## Methods

### Cell culture

The 4T1 murine breast cancer cell line was kindly provided by Dr Fred R. Miller (Michigan Cancer Foundation, Detroit, MI, USA). 4T1 cells were cultured in high glucose DMEM supplemented with 10% heat-inactivated FBS, 1% penicillin–streptomycin (P/S, from Gibco) and 1% Non-Essential Amino Acid (Gibco).

### Tumor models

4T1, MR13, sorted 4T1-Sca-1^+^ and 4T1-Sca-1^-^ (5×10^4^ cells in 50μl PBS/10% of 8.1 mg/ml Matrigel Matrix, were injected in the fourth right mammary gland of mice. Prior to surgery, ketamine (1.5 mg/kg) and xylazine (150 mg/kg) (both from Graeub) were injected intra-peritoneally to anesthetize the animals. Immune cell populations were analyzed at different time points post-tumor cell injection. Tumor length and width were measured twice a week with caliper and used to calculate tumor volume by the following equation: volume = (length x width^2^) x π/6. Tumors were collected and weighted at necropsy. For the intravenous injections, 2×10^5^ sorted 4T1-Sca-1^+^ and 4T1-Sca-1^-^ tumor cells resuspended in a volume of 50 μl of PBS were injected into the mice tail vein. Lung metastases were quantified 10 days post-injection. At each indicated time point, mice were sacrificed according to defined ethical criteria and were killed by CO_2_ inhalation followed by neck dislocation or terminal bleeding. All animal procedures were performed in accordance with the Swiss legislations on animal experimentation and approved by the Cantonal Veterinary Service of the Cantons Vaud and Fribourg for experiments in Lausanne and Fribourg (VD_1486.2; 2017_34_FR; 2017_34_FR, 2014_58_FR; 2011-33-FR).

### Reagents and chemicals

Growth factor reduced Matrigel Matrix (MG) was obtained from Becton Dickinson (BD Biosciences). Collagenase I was purchased from Worthington and DNAse I from Roche. Bovine serum albumin (BSA), crystal violet (CV) and paraformaldehyde (PFA) were obtained from Sigma-Aldrich. Drugs, inhibitors and cytokines: Doxorubicin and Methotrexate (generously provided by the Department of Oncology, University Hospital, University of Lausanne, Lausanne, Switzerland), Ruxolitinib (JAK inhibitor, Cat N°7064, Biotechne), anti-mouse Oncostatin M (R&D systems), anti-mouse IL-6 (BioxCell), recombinant mouse Oncostatin M and IL-6 (Biolegend, Cat #: 762802 & 575702 respectively).

### Antibodies

The following anti-mouse antibodies were used following manufacturer’s instructions: anti-CD16/CD32 Fc blocking antibody (BD Biosciences), anti-CD24-FITC (clone M1/69, eBioscience), anti-CD29-PE and-PE-Cy5 (clone HMβ1-1, BioLegend), anti-Sca-1-APC (clone D7, eBioscience), anti-CD61-Alexa 647 (clone 2C9.62(HMβ3-1), BioLegend, anti-CD45-PE (clone 30-F11, BD Biosciences), anti-Gr1-eFlour450 (clone RB6-8C5,eBioscience), anti-Ly6C FITC (clone HK1.4, BioLegend), anti-Ly6G-APC (clone 1A8-Ly6g,eBioscience), anti-F4/80 PerCP/Cy5 (clone BM8, eBioscience), anti-CD11b-PE-Cy7 (clone M1/70,eBioscience), anti-CD11c-APC-eFluor780 (clone N418, eBioscience), anti-CD4-FITC (clone 6K1.5, eBioscience), anti-CD8-PE (clone 53-6.7, eBioscience), anti-B220-APC (clone RA3-6B2, eBioscience), anti-CD49b-eFlour450 (clone DX5, BioLegend), Annexin V-APC (clone B217656, BioLegend), Propidium Iodide-PerCP (Clone V13245, Life technologies).

### Magnetic cell sorting (MACS)

MACS separators were used for positive and negative cell selections based on manufacturer’s instructions. Briefly, cells were counted and resuspend in 500 μl of MACS buffer with 10 μl of fluorescent coupled antibody of interest (APC-conjugated anti-Sca-1 and PE-conjugated anti-Gr1) per 10^7^ cells added. Cells were incubated for 30 minutes in the dark at 4°C and then washed with MACS buffer. After centrifugation and resuspension in 80 μl of MACS buffer per 10^7^ cells, 20 μl of anti-APC-conjugated magnetic beads (Miltenyi Biotec) per 10^7^ cells was added. After a 20 minutes incubation in the dark at 4°C, cells were washed with MACS buffer. After the centrifugation they were resuspended in MACS buffer and the magnetic separation was performed using LS MACS column (maximum 10^8^ labeled cells) for positive selection and LD MACS column (maximum 10^8^ labeled cells) for negative selection (Miltenyi Biotec). The purity of positive subpopulation was >70% and <99% for the negative subpopulation.

### Co-culture of Gr1^+^CD11b^+^ cells sorted from tumors or spleen of 4T1 tumor bearing mice

Gr1^+^ cells were sorted from tumors or spleen at day 23 post-injection (see below Magnetic beads cell sorting section). Afterwards, 1.5×10^5^ 4T1 cells were co-cultured in 6 wells plates using Transwell plates (0.4 μm, Nunc, ThermoFisher Scientific) with Gr1^+^CD11b^+^ sorted cells (top) and 4T1 cells (bottom) at different 4T1: Gr1^+^CD11b^+^ ratios (1:1, 1:3 and 1:5). After 48 hours of co-culture, cells were analyzed for Sca-1 expression and gene expression by flow cytometry and semi-quantitative real-time qPCR as indicated.

### Flow cytometry analysis on tissue samples

Mice were sacrificed at different time points for blood and tumors collection. Tumors were cut in small pieces with scissors, washed, and digested in serum free medium supplemented with Collagenase I and DNAse I (Roche). The mixture was incubated at 37 °C for 45 minutes on a shaking platform. Subsequently, serum-supplemented medium was added to neutralize the enzymatic reaction and the tissue suspensions were filtered through a 100 µm and a 70 µm sterile nylon gauzes. Upon centrifugation (5 minutes at 1400 rpm), pellets were recovered and red blood cells lysed with ACK buffer (Biolegend). The staining procedure and the flow cytometry acquisition are as described previously (81). Data acquisition was performed using the FACSCalibur (BD Biosciences) or MACSQuant flow cytometer from Miltenyi Biotec and data analyzed by FlowJo v10.0.7 (tree Stat Inc.).

### *In vitro* cell proliferation assay

Cells were collected and seeded in tissue culture 96-wells-plates (Costar) at 3’000 cells/well. Cells were grown in complete medium for 24, 36, 72 and 96 hours. At each time point cells were washed once with PBS, then fixed with 4% PFA and stained with 0.5% crystal violet solution for 0.5 hours. The stained cells were gently washed with deionized water to remove the extra dye and air-dried overnight at room temperature. After solubilizing the dye with crystal violet eluting buffer (70% ethanol and 1% acetic acid), cell viability was assessed by reading the absorbance at 595 nm wavelength in a multiwell plate reader (Modulus II microplate reader, Turner Biosystems). Results were analyzed by Prism (Graph pad software, Inc.) expressed as mean values of optical density (OD) of octuplet determinations ± SEM.

### *In vitro* cytotoxic assay

Tumor cells were plated at a concentration of 3’000 cell/well into 96-wells plates. The following day, a series of concentrations of the different drugs were supplemented to the culture medium. Untreated control cells were kept in normal culture medium. Cell viability of each well was assessed with crystal violet staining 48 hours after treatment, as described above. Results were analyzed by Prism software by a non-linear regression analysis and expressed as relative cell viability compared with non-treated control. The 50% maximum inhibition concentrations (IC_50_) were used to determine the drug-resistant ability of treated cells.

### Real-time reverse transcription qPCR and primers

Changes in mRNA expression levels were determined by semi-quantitative real-time qPCR. RNA samples were obtained from adherent cells using RNeasy kit from QIAGEN according to manufacturer’s instructions. From each sample, 1 μg RNA was retro-transcribed using SuperScript II Reverse Transcriptase kit (Life Technologies – Invitrogen), according to manufacturer’s instructions. The reactions were performed in a StepOnePlus™ thermocycler (Applied Biosystems, Life Technologies – Invitrogen) using the KapaSYBR® FAST SYBR Green Master Mix (Kapa Biosystems). Each reaction was performed in triplicates and values were normalized to murine 36B4 housekeeping gene. The comparative C_t_ method was used to calculate the difference of gene expression between samples. The following murine-specific primers (Microsynth AG) were used:

Sca-1 (F:5’-TCAGGAGGCAGCAGTTATTGTG-3’,R: 5’-TGGCAACAGGAAGTCTTCACG-3’), 36B4(F:5’-GTGTGTCTGCAGATCGGGTAC-3’,R:5’-CAGATGGATCAGCCAGGAAG-3’), OSM (F: 5’-ATGCAGACACGGCTTCTAAGA-3’, R: 5’-TTGGAGCAGCCACGATTG G-3’), OSMR: (F: 5’-CATCCCGAAGCGAAGTCTTGG-3’, R: 5’-GGCTGGGACAGTCCATTC TAAA-3’), IL-6: (F: 5’-TACCACTTCACAAGTCGGAGGC-3’, R: 5’-CTGCAAGTGCATCAT CGTTGTTC-3’).

### Histopathology

Tumors and lungs were harvested at the end of the experiments, fixed in formalin and embedded in paraffin. 5 μm thick serial sections were cut from the tissue blocks. 3-4 sections taken at 100 µm distance were stained with hematoxylin and eosin (H&E) and used to assess tumor morphology and quantify lung metastasis. Slides were scanned by Nanozoomer (Hamamatsu Photonics) and metastasis were counted manually using NDP.viewer2 software (Hamamatsu Photonics). Metastatic index was calculated by normalizing the metastasis number with the volume of primary tumor.

### Mammosphere-forming assay

5’000 cells/well were seeded in non-adhesive U bottom 96-wells plate in a semi-solid MEGM medium supplemented with 20 ng/ml EGF, and 20 ng/ml bFGF and heparin. Medium was gently replaced every 3 to 4 days. After 11 to 14 days culture, mammospheres of 50 to 150 μm diameter were detected under the microscope (bright field) and counted to quantify the sphere formation efficiency (SFE) as percentage of the initial number of seeded cells per well.

### Bulk RNA sequencing and data analysis

Four independent sorts of 4T1-Sca-1^+^ and 4T1-Sca-1^-^ cells (as described in the magnetic cell sorting section) or four independent 4T1 cells primed with Tu-Gr1^+^CD11b^+^ or Spl-Gr1^+^CD11b^+^ were prepared. RNAs of these cells were isolated using the NucleoSpin RNA protocol of Macherey-Nagel (as described in real-time (RT) qPCR and primers section). Samples were normalized for 1 μg RNA in a volume of 20 μl, sequenced on the NextSeq500 sequencer using the NextSeq 500/550 HT reagent v2 kit (Illumina) at the Swiss Integrative Center for Human Health (SICHH) in Fribourg, or the Lausanne Genomics Technologies Facility (GTF, UNIL) in Lausanne, Switzerland. For data analysis, all sequencing reads were processed for quality control, removal of low quality reads, adaptor sequence and ribosomal RNA by fastqc(0.11.8) (82), multiqc (1.9) (83), Trimmomatic (0.39) (84) and SortMeRNA(2.1) (85) accordingly. The filtered reads were mapped to the reference genome (mm10) using htseq-count (0.6.1) (86) or Salmon (0.99.0) (87). The normalization of the read counts and the analysis of the differential expression between the groups of samples were performed with in R(v4.1.3), a free software environment available at https://www.r-project.org/ using packages DESeq2 (v3.15) (88). Pathway enrichment analysis was performed using packages GSVA(1.42.0) (89) and GSEABase(1.56.0) (90) with the input of DESeq2 normalized count numbers using ssgsea method comparing the Hallmark genesets from MSigDB (v7.4.1) (91) with default settings. The significant altered pathways were determined by computing moderated t-statistics and false discovery rates with the limma(3.50.3)(92) for pair-wised comparison. The heatmaps were produced with R package pheatmap(1.0.12)(93) with default settings while pathway hierarchy clustering was performed by similarity based on Euclidean distance and the ward aggregation algorithm. The Sca1 Positive and Sca1 Negative signatures were extracted from the top 200 most upregulated or downregulated genes in 4T1-Sca-1^+^ and 4T1-Sca-1^-^ RNAseq data, respectively, with the threshold of adjusted p-value <0.05, fold change >1.5 or <-1.5 and average normalized count number >20. For Venn diagram, the genes fulfilling the threshold of adjusted p-value < 0.05, fold change > 1.5 or < -1.5 and average normalized count number >20 are compared. The figures were produced with R package venn (1.10) (94) . Further analysis and figures generation were performed in R using packages tidyverse(1.3.1)(95), ggplot2(3.3.6) (96), circlize(0.4.15) (97), biomaRt(2.50.3) (98, 99), RColorBrewer (1.1-3) (100), clusterProfiler (4.2.2) (101, 102), enrichplot (1.14.2) (103), ggpubr(v0.4.0)(104) ggbreak (0.1.0) (104).

### Microarray hybridization and data analysis

Experiments were performed as previously described (105). Briefly, triplicates wells of cultured 4T1 and MR13 cells were used for RNA extraction using RNeasy kit (QIAGEN). Probe synthesis and GeneChip Mouse Gene Exon 1.0 ST Array (Affymetrix Ltd) hybridization were performed at the GTF, UNIL, Lausanne, Switzerland. Microarray analyses were carried out with R. After quantification of gene expression with robust multi-array normalization (99) using the BioConductor package Affy, (http://www.bioconductor.org/) significance of differential gene expression was determined by computing moderated t-statistics and false discovery rates with the limma package (92). Annotation was based on the genome version NCBI Build 36 (Feb. 2006). The obtained p-values were corrected for multiple testing by calculating estimated false discovery rates (FDR) using the method of Benjamini-Hochberg. Heatmaps were produced by color-coding gene-wise standardized log gene expression levels (mean zero standard deviation one). Probe-sets were shown hierarchically clustered by similarity based on Euclidean distance and the ward aggregation algorithm.

### Public RNAseq data analysis

The RPKM normalized gene expression data in Ross dataset (GSE150928) from multiple murine models of breast cancer metastasis was obtained from Gene Expression Omnibus (GEO) in the NCBI data repository. The data were analyzed and plot with R package tidyverse(1.3.1)(95), ggplot2 (3.3.6) (96) and ggpubr (v0.4.0) (106). The single-cell RNAseq data in Sebastian dataset (43) was obtained from Dryad data repository (https://doi.org/10.6071/M3238R). The data were analyzed with R package Seurat (3.0) (107). The tumor and different myeloid cell populations were extracted for cell-cell interaction analysis using CellPhondDB (2.0) (44). The identified interaction pairs were extracted and plot using circlize (0.4.15) (97) and ComplexHeatmap (2.11.2) (108). The ligands and receptors annotated in the circular plot were compiled from databases in CellTalkDB(109), SingleCellSignalR (110). To investigate the tumor cell dynamics, datasets for 4T1 (GSE158844 and GSM3502134) and MCF-7 (GSM4681765 and GSM5904917) were obtained from GEO. The data were filtered and normalized separately before merging with IntegrateData function included in Seurat. Cell cycle regression was performed according to the standard protocol of Seurat. For MCF-7, due the huge difference of the sample size, 2000 cells were randomly selected from GSM4681765 data, and then merged with GSM5904917. Single-cell trajectories analysis was performed with monocle 3 (111–114) with default settings and the root and start point were selected manually for pseudotime caculation. The GSEA analysis of selected clusters was performed with R package fgsea(115) and the gene rank was calculated with Wilcoxon rank sum test and auROC analysis using wilcoxauc function included in presto package. The Tu-Gr1^+^CD11b^+^-induced signature was extracted by comparing gene expression between Tu-Gr1^+^CD11b^+^ and Spl-Gr1^+^CD11b^+^-educated 4T1 cells with adjusted p-value <0.05, fold change >2, and the top 50 genes were selected.

### Clinical data analysis

To validate our finding in clinical data, the human orthologs of murine Tu-Gr1^+^CD11b^+^-induced signature genes were used. Conversion from murine to human gene symbols and Entrez IDs was performed with the biomaRt package (2.46.3) (98, 99), using the reference mart https://dec2021.archive.ensembl.org. Molecular Taxonomy of Breast Cancer International Consortium (METABRIC) breast cancer data was downloaded from cBioPortal (116–118) in August 2022, and expression data was log2 transformed. Expression values were stratified in two groups by median values. Survival curves were generated using the ggsurvplot function from the survminer package (0.4.9)(106), and were compared between groups using a log-rank test. Survival curves were created using the survfit function from the survival package (3.2.11) (119). Cox proportional hazard regression model was performed through the coxph function of the same package.

### Statistical analyses

Unless specified, the data were presented as mean ± SEM from at least 3 independent experiments, unless otherwise indicated. Statistical comparisons were performed by an unpaired Student’s t test with a two-tailed distribution, one-way ANOVA analysis of variance with Bonferroni post-test or two-way ANOVA with Tukey’s or Dunnett’s multiple comparison test using Prism 7.0 GraphPad Software, Inc. or R package ggpubr (v0.4.0)(104).

### Graphic illustrations

Illustrative schemes were created with BioRender.com.

### Data and code availability

The raw and processed bulk RNAseq data used to generate figures in Figure 3 and Supplemental Figure 5 and microarray data used to generate figures in Figure 7 have been deposited in the GEO database under the access code GSEXXXXXX. The code used for the analyses is open-source and available through with R packages described in methods.

## Author contributions

Conceptualization, S.P., Q.L., C.R.; Methodology, S.P., N.F., Q.L., C.R.; Software, A.K., N.F., Q.L.; Validation, S.P., M.B., A.K., Q.L.; Formal analysis, S.P., M.B., A.K., Q.L.; Investigation, S.P., M.B., A.K., O.C., Y-T.H., N.D., L.G., Q.L.; Resources, S.P., M.B., A.K., O.C., N.D., L.G., G.L.; Data Curation, S.P., M.B., A.K., N.D., Q.L., C.R.; Writing – original draft preparation, S.P., Q.L., C.R.; Writing – review and editing, S.P., M.B., A.K., Y-T.H., G.L., N.D., Q.L., C.R.; Visualization, S.P., M.B., A.K., Q.L., C.R.; Supervision, S.P., Q.L., C.R.; Project administration, Q.L., C.R.; Funding acquisition, C.R.

## Acknowledgments

The authors wish to thank Sarah Cattin and Melissa Rizzo for assistance with FACS analysis, Dr. Ana-Marija Sulić (Institute of Biotechnology, HiLIFE, University of Helsinki) for the insightful discussion about scRNA-seq data analysis, Dr. Fred R. Miller (Michigan Cancer Foundation, Detroit,MI, USA) for providing 4T1 cells, Dr. Khalil Zaman (Department of Oncology, University Hospital, University of Lausanne, Lausanne, Switzerland) for providing chemotherapeutic drugs. This work was supported by grants from the Swiss National Science Foundation (31003A_179248; 310030_208136), the Swiss Cancer League (KFS 4400-02-2018) and the Medic Foundation (to C.R.).

