## Supplemental Figures for "Tumor-educated Gr1^+^CD11b^+^ cells instigate breast cancer metastasis by twisting cancer cells plasticity via OSM/IL6–JAK signaling"

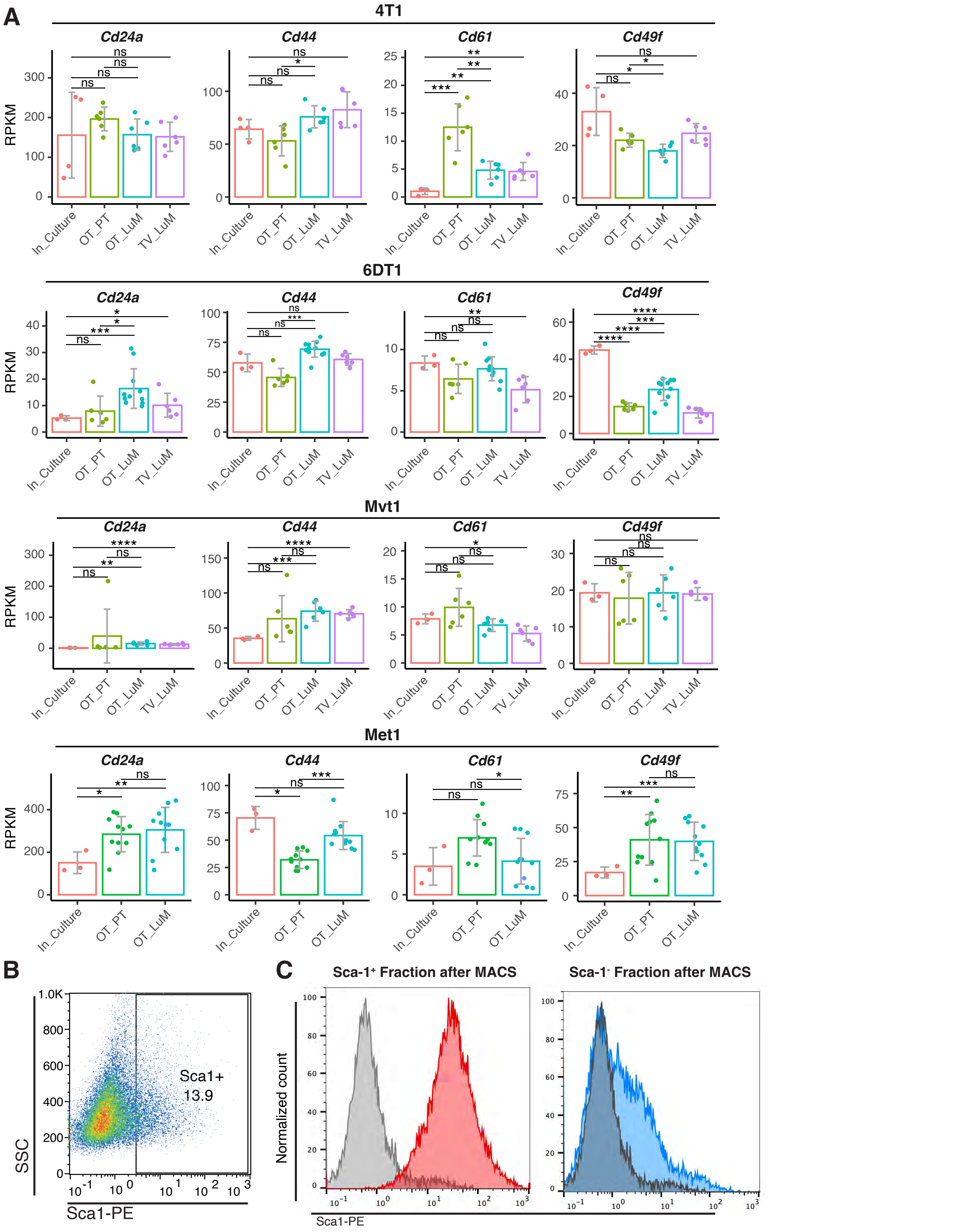


**Supplemental Figure 1. Stem cell markers expression in multiple metastatic breast cancer models**
**(A)** Stem cell marker *Cd24a*, *Cd44*, *Cd61* and *Cd49e* mRNA expression in 4T1, 6DT1, Mvt1 and Met1 metastatic murine breast cancer models extracted from Ross dataset. Data are presented as mean values of RPKM ± SD. *, p<0.05; **, p< 0.01; ***, p< 0.001; ****, p< 0.0001; ns, non-significant (unpaired two-tailed student's t test with Holm correction).
**(B)** Dot plot representation of Sca-1 expression in 4T1 parental cell line determined by flow cytometry.
**(C)** Histogram of Sca-1 expression distribution on MACS positively selected Sca-1$^{+}$ cells (red histogram) and MACS negatively selected Sca-1$^{-}$ cell (blue histogram). Gray: histogram of fluorescence of unstained cells.

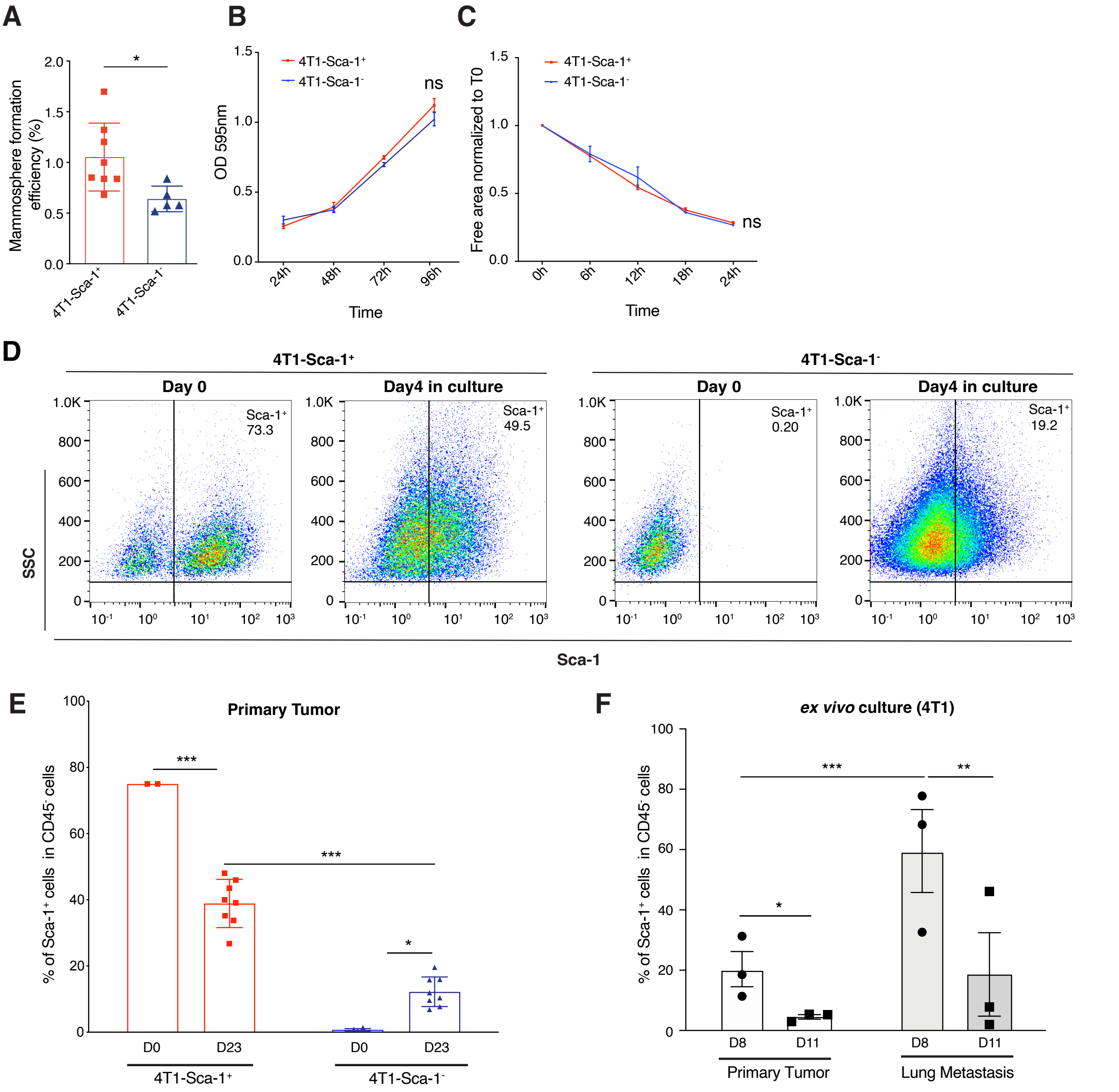


**Supplemental Figure 2. Sca-1⁺ tumor cells have stem cell-like features in vitro and show plasticity in vivo and in vitro**
**(A)** Quantification of the mammosphere forming efficiency of 4T1-Sca-1⁺ and 4T1-Sca-1⁻ populations (n=5-8/group).
**(B)** Cell proliferation curve of 4T1-Sca-1⁺ and 4T1-Sca-1⁻ in vitro MACS isolated tumor cells determined by crystal violet assay. The results represent optical density (OD) of the wells (n=5-6/group).
**(C)** Cell motility of 4T1-Sca-1⁺ and 4T1-Sca-1⁻ sorted tumor cells determined by a scratch wound healing assay (n=5-6/group). Results are given cell-free area relative to the initial wound area.
**(D)** The abundance of Sca-1⁺ population after MACS isolation from parental 4T1 cells. Day 0: immediately after positive sorting, the Sca-1⁺ population to accounts for 73.3% of total cells (upper panel). After 4 days of in vitro culture the abundance of the Sca-1⁺ population decreased to 48.9%. Negatively sorted Sca-1⁻ population accounts for >99 % of total cells (lower panel). After 4 days of in vitro culture 19.2% of the initially Sca-1⁺ cells were Sca-1⁺.
**(E)** Abundance of Sca-1⁺ population at the time of orthotopic injection of MACS isolated 4T1-Sca-1⁺ and 4T1-Sca-1⁻ cells into the mammary fat pad of BALB/C mice (D0) and in the derived primary tumors 23 days post injection (D23) as indicated. n=3-8/group.
**(F)** Abundancy of the Sca-1⁺ population in tumor cells recovered from primary tumors and lung metastases 21 days post orthotropic injection and further cultured for 8- and 11-days ex vivo as indicated. n=3/group.
Data represent mean values ± SEM from 3 independent experiments. P values were calculated using unpaired two-tailed student's t test **(A)**, or two-way ANOVA with Tukey multiple-comparison test **(B, C)**, and one-way ANOVA with Tukey multiple-comparison test **(E, F)**. *, p<0.05; **, p< 0.01; ***, p< 0.001, ns, non-significant.

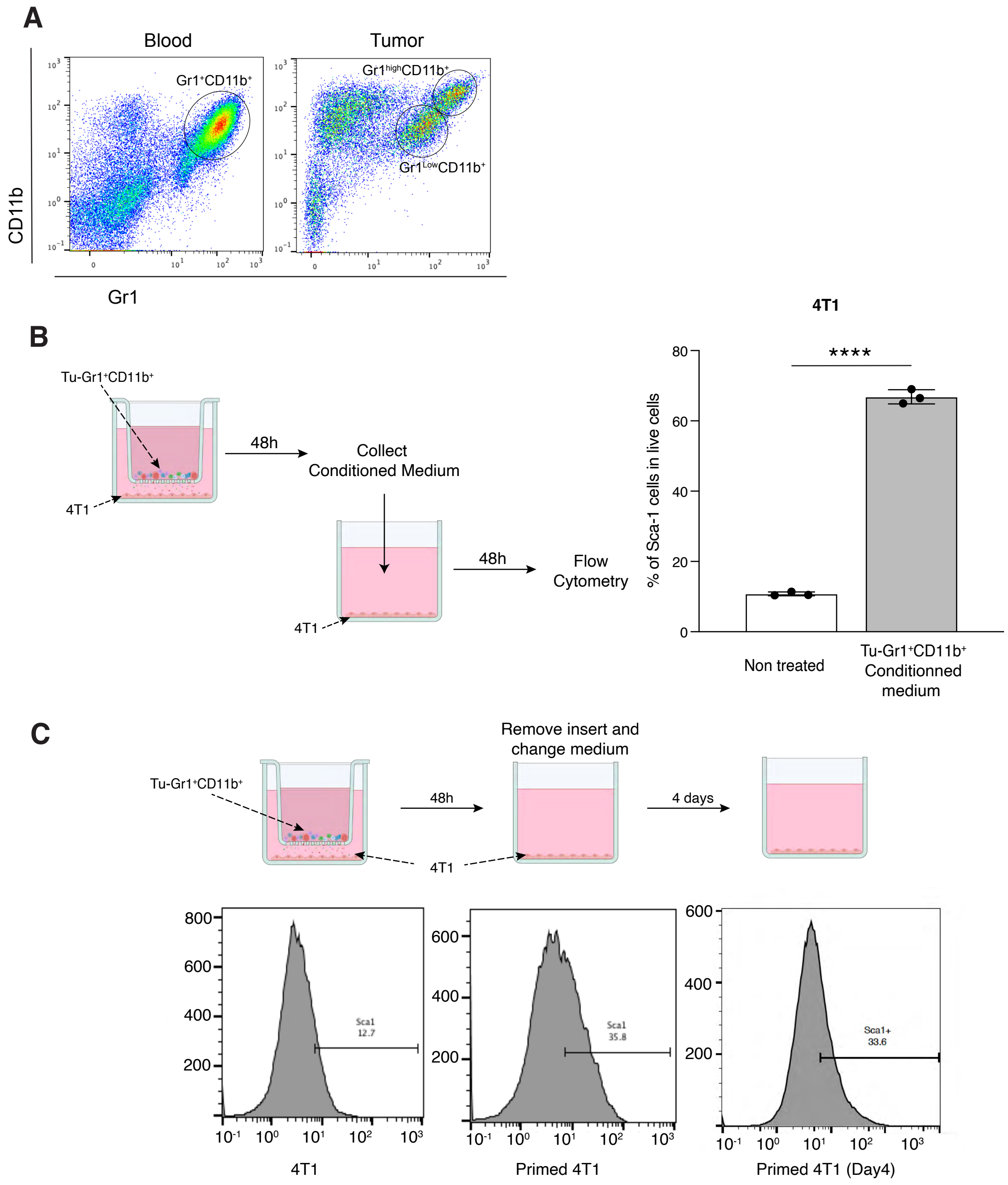


**Supplemental Figure 3. Tu-Gr1⁺CD11b⁺-derived secreted factors promote the enrichment of Sca-1⁺ population**

**(A)** Dot plots presenting Gr1⁺CD11b⁺ subpopulations in Blood and Tumor site of 4T1 tumor-bearing mice.

**(B)** Abundance of the Sca-1⁺ population in tumor cells cultured for 48 hours in the presence of medium collected from 4T1/Tu-Gr1⁺CD11b⁺ cocultures (48 hours). Sca-1 expression on 4T1 cells was determined by flow cytometry (n = 3/group). Data represent mean values ± SEM from 3 independent experiments. ****, p< 0.0001 (unpaired two-tailed student's t test).

**(C)** Sca-1 expression in parental 4T1 cells (left panel), 4T1 primed for 2 days with Gr1⁺CD11b⁺ cells (middle panel) and cultured for 4 additional days in the absence of Gr1⁺CD11b⁺ (right panel). Sca-1 expression was determined by flow cytometry.

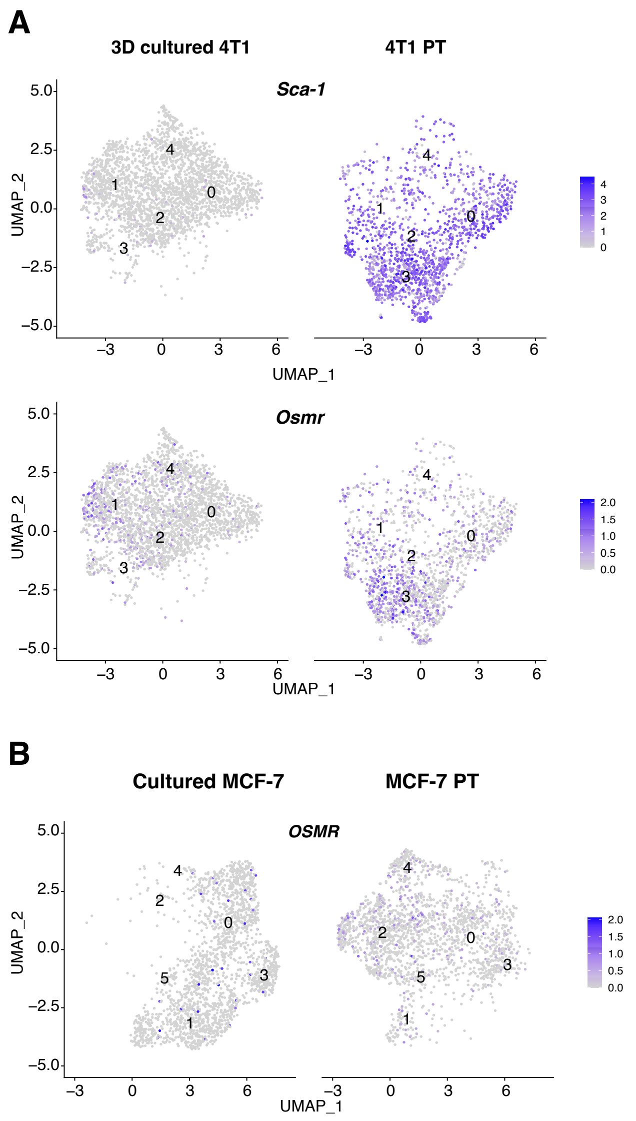


**Supplemental Figure 4. TME expands the Sca1⁺ population during in vivo tumor progression**
(A) UMAP plots showing the expression of *Sca-1* and *Osmr* in 4T1 tumor cells in 3D culture or primary tumors (PT). Analysis based on publicly available data (GSM4812003 and GSM3502134)
(B) UMAP plot showing the expression of *OSMR* in MCF-7 tumor cells in culture or primary tumors (PT). Analysis based on publicly available data (GSM4681765 and GSM5904917).

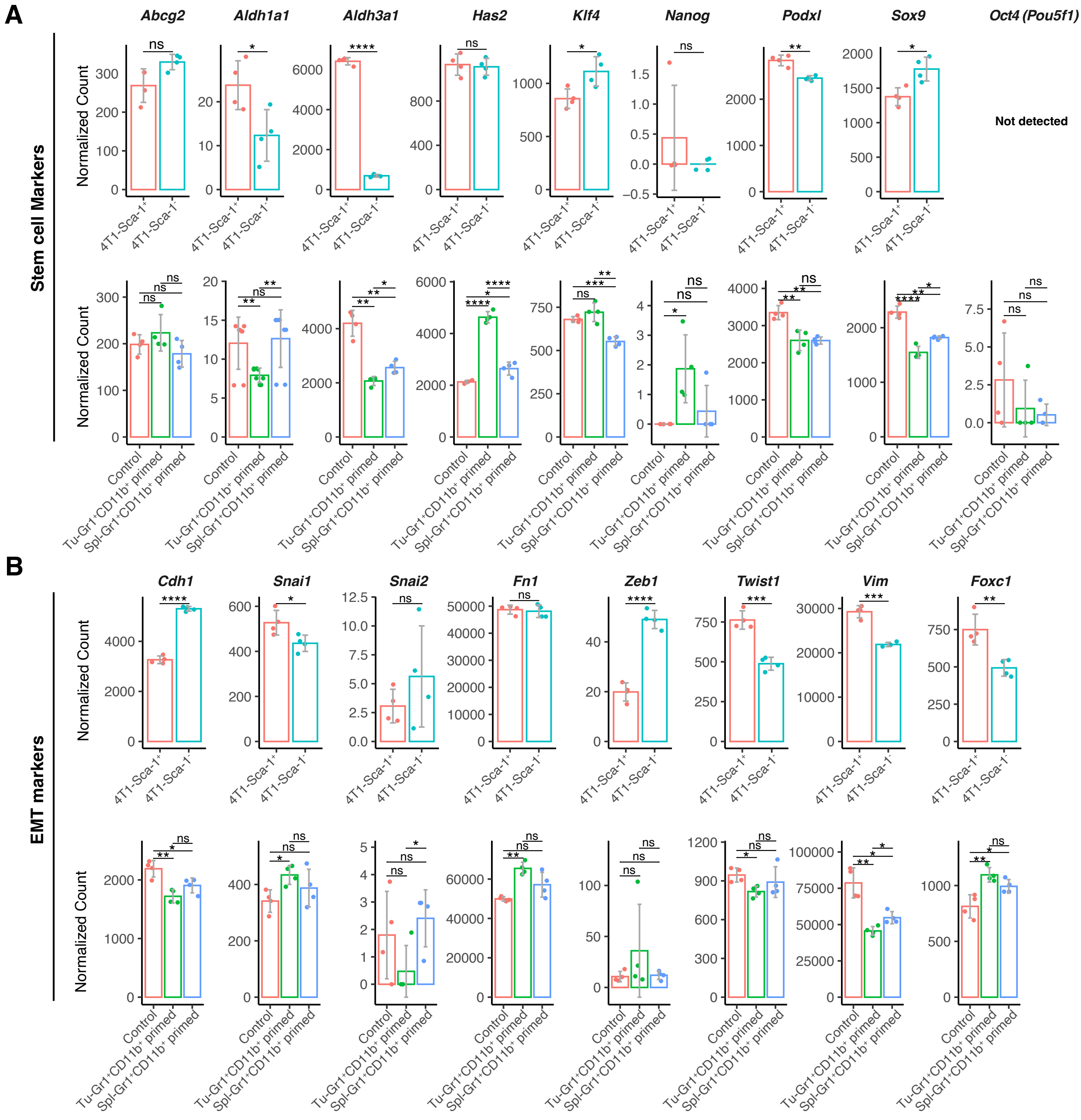


**Supplemental Figure 5. Expression of selected stem cell/CSC and EMT genes in 4T1-Sca-1$^{+}$, 4T1-Sca-1$^{-}$ and differently primed 4T1 cells**

Expression of stem cell markers **(A)** and EMT markers **(B)** in 4T1-Sca-1$^{+}$ cells, 4T1-Sca-1$^{-}$ cells, and parental, Tu-Gr1$^{+}$CD11b$^{+}$-educated, Spl-Gr1$^{+}$CD11b$^{+}$-educated 4T1 cells, as indicated.

Data represent mean values of RPKM ± SD. *, p<0.05; **, p< 0.01; ***, p< 0.001; ****, p< 0.0001 (unpaired two-tailed student's t test, and with Holm correction for Gr1$^{+}$CD11b$^{+}$-educated 4T1 cells).

A

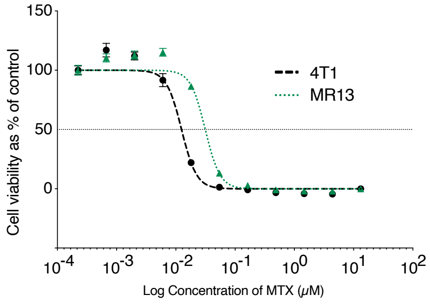


B

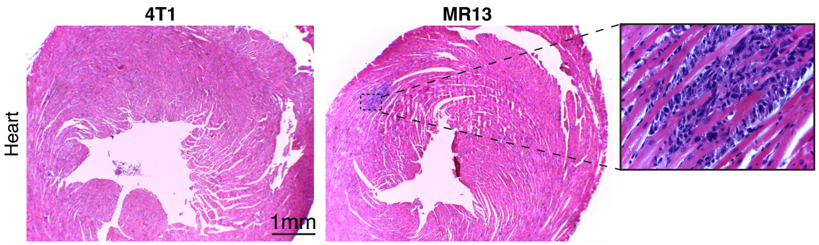


**Supplemental Figure 6. Sca-1⁺ enriched 4T1 cells and MR13 cells are more resistant to chemotherapy drugs**
**(A)** Dose dependent effect of 48 hours treatment with MTX on 4T1 and MR13 cells' viability. IC50 of MTX for 4T1 and MR13 are 12.5 nM and 31.4 nM respectively.
**(B)** Illustrative H&E stained heart sections of BALB/c mice 22 days post orthotopic injection with 4T1 cells (left) or MR13 cells (right and insert) (n=8-9/group).

**A**

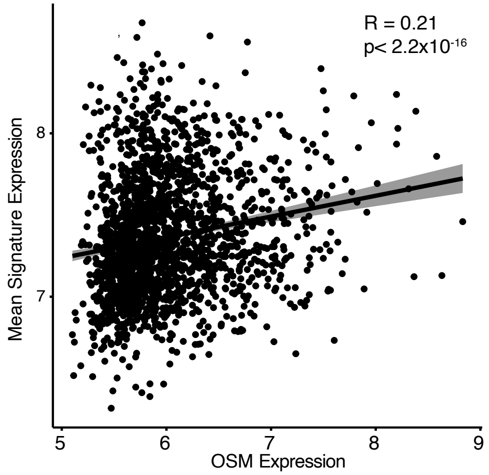


**Supplemental Figure 7. Genes of the Tu-Gr1⁺CD11b⁺-induced signature and patients' outcome**
(A) Correlation of the orthologue 32 gene signature, based on the Tu-Gr1⁺CD11b⁺-induced signature, with OSM expression in the METABRIC datasets. Spearman's correlation coefficients and p-values are shown.

**Supplemental Table 1.  Gene signature related to Figure 3**

**Supplemental Table 2. CellPhoneDB analysis results**

**Supplemental Table 3. Tu-Gr1$^{+}$CD11b$^{+}$-induced signature**
